# Mapping acute neuroinflammation *in vivo* with diffusion-MRI in rats given a systemic lipopolysaccharide challenge

**DOI:** 10.1101/2022.11.22.517484

**Authors:** Eugene Kim, Ines Carreira Figueiredo, Camilla Simmons, Karen Randall, Loreto Rojo Gonzalez, Tobias Wood, Brigida Ranieri, Paula Sureda-Gibert, Oliver Howes, Carmine Pariante, NIMA Consortium, Ofer Pasternak, Flavio Dell’Acqua, Federico Turkheimer, Diana Cash

## Abstract

It is becoming increasingly apparent that neuroinflammation plays a critical role in an array of neurological and psychiatric disorders. Recent studies have demonstrated the potential of diffusion MRI (dMRI) to characterize changes in microglial density and morphology associated with neuroinflammation, but these were conducted mostly *ex vivo* and/or in extreme, non-physiological animal models. Here, we build upon these studies by investigating the utility of well-established dMRI methods to detect neuroinflammation *in vivo* in a more clinically relevant animal model of sickness behavior. We show that diffusion tensor imaging (DTI) and neurite orientation dispersion and density imaging (NODDI) indicate widespread increases in diffusivity in the brains of rats given a systemic lipopolysaccharide challenge (n=20) vs. vehicle-treated controls (n=12). These diffusivity changes correlated with histologically measured changes in microglial morphology, confirming the sensitivity of dMRI to neuroinflammatory processes. This study marks a further step towards establishing a noninvasive indicator of neuroinflammation, which would greatly facilitate early diagnosis and treatment monitoring in various neurological and psychiatric diseases.

## 1 Introduction

In recent years, evidence has accumulated to suggest that neuroinflammation plays a crucial role in a wide range of neurological and psychiatric disorders including depression and mood disorders (Kwon and Koh, 2020; Miller et al., 2017; Mondelli et al., 2017; Nettis and Pariante, 2020). There is an intricate balance between the beneficial effects of an operating immune system, such as euflammation and protective sickness behavior, and the adverse consequences of chronic inflammation that can potentially lead to neural damage (Pape et al., 2019). Neuroinflammation is largely mediated by reactive microglia, the central nervous system’s primary immune cells, and is characterized by a wide and varied network of local and distant responses based on the type, intensity, and duration of a given trigger. An example is increased blood-brain barrier (BBB) permeability which can play a role in neuroinflammation as witnessed in active focal lesions in multiple sclerosis (Gaitan et al., 2011) and in Alzheimer’s Disease (AD) (Bowman and Quinn, 2008) although its occurrence in other disorders such as lupus and depression was recently challenged (Althubaity et al., 2022; Stock et al., 2017; Turkheimer et al., 2022; Turkheimer et al., 2021).

Detecting neuroinflammation in living humans would be of great benefit to early disease detection and monitoring treatment efficacy, particularly in diseases that lack robust pathological hallmarks such as psychosis and mood disorders. Although this is not yet reliably possible, an array of nuclear imaging and magnetic resonance (MR)-based methods can identify and characterize neuroinflammatory processes *in vivo* (Albrecht et al., 2016). The current gold-standard is positron emission tomography (PET) using tracers that target the 18kDa translocator protein (TSPO) (Van Camp et al., 2021; Zhang et al., 2021). TSPO is found in the mitochondrial membranes and mostly expressed in endothelial cells, astrocytes, and microglia; its upregulation in microglia, infiltrating macrophages and, to a lesser extent, astrocytes is generally associated with neuroinflammatory events (Guilarte et al., 2022). However, although widely seen as the best available neuroinflammation biomarker, it presents many limitations. These include substantial within- and between-subject variability in TSPO radioligand binding (Collste et al., 2016) and restriction to small sample sizes given the high cost, logistical burden, ionizing radiation, and availability of the technique. These factors limit the utility of TSPO as a marker of inflammation, further complicated by a lack of clarity about its exact cellular sources and mechanism of action in this context (Notter et al., 2021; Nutma et al., 2022; Vicente-Rodriguez et al., 2021).

Although MR methods are overall limited by a lack of specificity when compared to nuclear imaging, they are free of ionizing radiation and able to measure multiple tissue characteristics (Albrecht et al., 2016). Magnetic resonance spectroscopy (MRS) measures metabolite concentrations such as N-acetyl-aspartate (NAA), myo-inositol and choline compounds, making it a valuable method to identify glial cell changes after their migration to or activation in the inflamed area (Chang et al., 2013). Dynamic contrast-enhanced MRI can detect BBB disruption by measuring the leakage of gadolinium-based contrast agents (Heye et al., 2014), while a recently developed method based on water exchange rather than contrast agent leakage has been used to detect subtle BBB breakdown in a rat model of AD (Dickie et al., 2019).

Diffusion-weighted magnetic resonance imaging (dMRI) is a suite of techniques that can characterize brain microstructure *in vivo* and identify subtle abnormalities (Karlsgodt, 2019) including those occurring during neuroinflammation. A type of dMRI, diffusion tensor imaging (DTI), has been reasonably successful in detecting neuroinflammation-related brain changes. For example, DTI has revealed increased axial diffusivity in the corpus callosum in rat models of sepsis-associated encephalopathy (Dhaya et al., 2018; Griton et al., 2020). Other studies have shown increased diffusivity to be a robust marker of neurodegenerative disorders associated with neuroinflammation (Agosta et al., 2011; Inglese and Bester, 2010), although this may reflect a number of different aspects such as BBB integrity, fiber density, myelination, tissue organization, and membrane permeability (Jones et al., 2013). In addition to lacking specificity, the DTI model and its processing tools were originally designed for white matter and axons (Klawiter et al., 2011) complicating the biological interpretation of DTI changes in the gray matter and parenchyma of the brain.

More recent and sophisticated diffusion models attempt to separate different tissue compartments. For example, the free-water imaging (FWI) model estimates the fractional volume of freely diffusing water (Pasternak et al., 2009), which has been found to be increased in both white and gray matter in first-episode schizophrenia patients and attributed to increased extracellular space due to neuroinflammation (Pasternak et al., 2012). The neurite orientation and dispersion density imaging (NODDI) model further separates the tissue into putative neurite, extra-neurite, and cerebrospinal fluid (CSF) compartments (Zhang et al., 2012). NODDI has been used to characterize gray matter microstructure (Nazeri et al., 2020) and, along with other advanced diffusion techniques, has been demonstrated to be sensitive to microglial density (Yi et al., 2019) and inflammation-induced changes in glial microstructure (Garcia-Hernandez et al., 2022).

Preclinical studies are essential for validation of most imaging biomarkers, and animal models of neuroinflammation play a central role in the search for the underlying biological mechanisms responsible for these changes. Some of the well-established rodent models of neuroinflammation use either intracranial or systemic administration of lipopolysaccharide (LPS), a component of the Gram-negative bacteria cell wall, to induce an immune response. This activates glial cells *in vivo*, causing a consequent inflammatory cascade that includes the release of inflammatory cytokines, chemokines, kynurenine, and nitric oxide (NO), all of which are thought to contribute to neurotoxicity (Dantzer et al., 2008; O’Connor et al., 2009). This has supported the LPS challenge of the immune system as a valuable model to mimic the effects of neuroinflammatory events and their consequences (Batista et al., 2019; Biesmans et al., 2013; Hoogland et al., 2015; Zhao et al., 2019).

With this in mind, we aimed to image neuroinflammation as indexed by microglial activation in response to systemic (intraperitoneal) administration of a low dose of bacterial LPS in a well-known rodent model of sickness behavior which has also been used to study aspects of mood disorders such as depression (Biesmans et al., 2013; Konsman et al., 2002). Because this approach induces mild neuroinflammation indirectly, as a response to a systemic inflammatory cascade, and does not feature invasive intracranial administration, this LPS model is considered more representative of the clinical scenario. Given the paucity of established *in vivo* biomarkers in this type of clinically relevant model, we explored three different dMRI models – DTI, FWI, and NODDI – to compare their distinct characteristics and sensitivities to regional inflammatory LPS-induced changes in the brain.

## 2 Materials and Methods

### 2.1 Animals

All animals were treated in accordance with the Animals (Scientific Procedures) Act 1986 and KCL local ethical guidelines. Adult male Sprague Dawley (Charles River, UK) rats weighing 220-240g were fed *ad libitum* by standard rat chow and tap water and housed in an airconditioned room with an ambient temperature of 21± 3°C and a 12-hour light/dark cycle (lights on at 7am and lights off at 7pm).

### 2.2 Experimental design

LPS (*E. coli*, O111:B4 serotype, Sigma Aldrich, UK) was dissolved at 0.5 mg/ml in sterile 0.9% saline. Rats were weighed and tested for locomotor behavior before being injected intraperitoneally (i.p.) with 0.5 mg/kg LPS (n=20) or 1 ml/kg saline vehicle (n=12) during the lights-on phase (i.e., their inactive period). Twenty-two (±1) hours later, they were weighed and tested for locomotor behavior and then scanned (24±1h after LPS) using MRI before being killed, and blood and brain samples harvested for biochemical and histological analyses.This dose of 0.5mg/kg LPS was chosen because it has been shown in published reports to reliably induce symptoms of sickness behavior associated with depression (Biesmans et al., 2013; Ilanges et al., 2022; O’Connor et al., 2009; Zhao et al., 2019). Furthermore, our in-house pilot data confirmed that this is the lowest dose (compared to 0.1 and 0.25 mg/kg) that can induce microglia proliferation under our experimental conditions.

### 2.3 Locomotor behavior

Animals were habituated to the open field arena for 30 minutes per day for four consecutive days 1-3 days before testing. The arena consisted of a 120×120-cm Perspex box divided into 60×60-cm quadrants. One animal was placed in each quadrant for 30 minutes (with up to four animals tested simultaneously). On the test day, animals were placed in the open field for 30 minutes and allowed to move freely. They were filmed using an overhead mounted camera and recorded using Open Broadcaster Software (https://obsproject.com/). Videos were then analyzed for time spent moving, velocity, and distance moved using TopScan Suite (CleverSys Inc., Reston, VA, USA).

### 2.4 MRI acquisition

After locomotor behavioral testing, the rats were imaged *in vivo* in a Bruker BioSpec 9.4T scanner (Bruker BioSpin MRI GmbH, Ettlingen, Germany) using an 86-mm volume resonator for transmission and a 4-channel surface array receive-only coil. During scanning, the rats were anesthetized with isoflurane (∼2% in medical air [1 L/min] + medical O_2_ [0.4 L/min]), which was adjusted as needed to keep the respiration rate between 50 and 60 breaths/min. The rats’ internal temperatures were monitored with a rectal probe and maintained at 37°C via a warm water circulation system (Small Animal Instruments Inc., Stony Brook, NY, USA).

For each rat, a T2-weighted (T2w) structural reference image and diffusion-weighted images were acquired using the following sequences:

- 3D RARE (Rapid Acquisition with Relaxation Enhancement): echo time (TE) = 32 ms, repetition time (TR) = 1800 ms, RARE factor = 12, field-of-view (FOV) = 30×20×20 mm, matrix = 192×128×64, scan time = 14 min 42 s.
- 2D single-shot pulsed-gradient spin-echo EPI (Echo Planar Imaging): TE/TR = 24.5/9000 ms; FOV = 21.12×17.16 mm; matrix = 64×52; 64 contiguous slices of 0.33-mm thickness; gradient duration (δ) = 4 ms; gradient separation (Δ) = 10 ms; b-values = 500, 1000, and 1500 s/mm^2^; 60 diffusion directions per shell; 6 b0 images; scan time ∼40 min with respiration gating. Six additional b0 images with opposite phase encoding polarity were acquired for correction of susceptibility-induced distortions.

### 2.5 MRI processing

Susceptibility- and eddy-current-induced distortion and motion correction were applied to the diffusion images using topup (Andersson et al., 2003; Smith et al., 2004) and eddy (Andersson and Sotiropoulos, 2016) from the FMRIB Software Library (FSL). Three different diffusion models were fit to the preprocessed data:

1. diffusion tensor imaging (**DTI**) model using FSL `dtifit`,
2. free water imaging (**FWI**) model using DIPY `fwdti`,
3. neurite orientation dispersion and density imaging (**NODDI**) model using the NODDI Matlab Toolbox.

All three diffusion shells (b500, b1000, and b1500) were used to fit the three models. In addition, DTI models were fit using each shell individually to assess potential effects in the slower, intermediate, and faster diffusion regimes separately. Group-level voxel-wise statistics were performed on the fractional anisotropy and mean diffusivity values calculated from these different DTI fits, which will be referred to as FA and MD (all three shells), FA500 and MD500 (b500 shell only), FA1000 and MD1000 (b1000 shell only), and FA1500 and MD1500 (b1500 shell only).

The DTI model consists of a single compartment for all tissue. The FWI model expands upon this by adding a second compartment in which water diffuses freely in an unhindered, isotropic manner (Pasternak et al., 2009), introducing an extra free parameter f, the volume fraction of the tissue compartment. Voxel-wise statistics were performed on the estimated volume fraction of the free water compartment, (1 – f), referred to here as FW, as well as on FA and MD of the tissue compartment (FAt and MDt).

Building on the FWI model, the NODDI model consists of three compartments originally described as the intra-cellular, extra-cellular, and cerebrospinal fluid (CSF) compartments (Zhang et al., 2012):

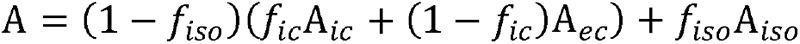

A is the MR signal attenuation of a diffusion-weighted image compared with a non-weighted b0 image, A_*ic*_ and f_*ic*_ are the signal attenuation and volume fraction of the intra-cellular compartment, A_*ec*_ and f_*ec*_ are the signal attenuation and volume fraction of the extra-cellular compartment, and A_*iso*_ and f_*iso*_ are the signal attenuation and volume fraction of the CSF compartment (i.e., the isotropic diffusion compartment).

These compartments can be renamed to be more biologically agnostic as the restricted diffusion, hindered diffusion, and unhindered isotropic diffusion compartments, respectively:

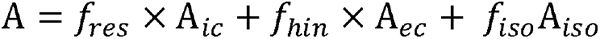

f_*res*_ – the restricted diffusion volume fraction – is defined as:

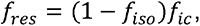

f_*hin*_ – the hindered diffusion volume fraction – is defined as:

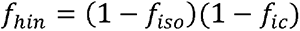

Voxel-wise statistics were performed on the three NODDI compartment volume fractions as well as the orientation dispersion index (ODI).

ANTs (Advanced Normalization Tools) was used to construct a multimodal study template from the T2w images and FA maps (`antsMultivariateTemplateConstruction2.sh`) (Avants et al., 2011). The FA map of each brain was normalized to the FA template via sequential rigid-body, affine, and diffeomorphic (SyN) (Avants et al., 2008) registrations using `antsRegistration`, and the resultant transformations were applied to the other DTI, FWI, and NODDI parametric maps. Group mean images of the parametric maps are shown in **Supplemental Figure 3**.

### 2.6 MRI analysis

Voxel-wise Spearman correlation coefficients were calculated across all animals to compare diffusion parameters computed from the three different models. Voxel-based analysis was performed using permutation tests and threshold-free cluster enhancement (TFCE) to test for differences in diffusion metrics between the LPS and vehicle control groups; p-values were corrected for multiple comparisons by controlling the family-wise error (FWE) rate (FSL `randomise`) (Winkler et al., 2014). Based on the results of the voxel-wise analysis, a subset of parameters (MD500, MD1500, FW, and f_*res*_) was chosen for further ROI-based analysis.

A hybrid rat brain atlas was made in-house by combining the Waxholm Space Sprague Dawley atlas of Papp et al. (Papp et al., 2014), which has detailed parcellation of subcortical regions but very few cortical regions, with the *in vivo* cortical atlas of Valdes-Hernandez et al. (Valdes-Hernandez et al., 2011), resulting in parcellation of the whole brain into 115 regions of interest (ROI). For this study, the number of ROIs was reduced to 64 by merging anatomical subregions, e.g., the lateral and medial regions of the parietal association cortex (**Supplemental Figure 1**). This new atlas was then non-linearly registered to the study template space. After down-sampling to the diffusion MRI resolution, five ROIs (alveus, central canal, fasciculus retroflexus, habenular commissure, and mammillothalamic tract) had volumes of fewer than five voxels and were excluded from all further analysis. The median values of MD500, MD1500, FW, and f_*res*_ were calculated for each of the remaining 59 ROIs for each rat. Mann-Whitney U tests were used to test for univariate differences between groups for each ROI, and the Benjamini-Hochberg step-up procedure used to control the false discovery (FDR) rate for each parameter separately using MATLAB (MathWorks, Natick, MA, USA).

### 2.7 Sample collection and plasma cytokine measurements

Immediately after scanning, animals were over-anaesthetized with isoflurane (5% in medical air [1 L/min] + medical O_2_ [0.4 L/min]) and decapitated. Trunk blood was taken and centrifuged to separate blood cells from plasma. Plasma was collected and stored at – 20°C and later analyzed for IL-6, IL-1b, TNF-a, and KC/GRO levels using a custom multi-spot immunoassay V-Plex kit (Meso Scale Discovery LLC, Rockville, USA), and for acute phase protein α-2-macroglobulin using an ELISA kit (Life Diagnostics Inc., West Chester, USA).

The brain was removed from the skull and divided in two hemispheres. One hemisphere was immersion fixed for 48 hours in 4% formaldehyde before being cryopreserved in 30% sucrose for histological analysis. The second hemisphere was divided in two sections along the midline, and each section was snap frozen in isopentane cooled to ca. -40°C. The medial section was later analyzed for cytokines using the same custom MSD immunoassay V-Plex kit mentioned above. Protein levels of the brain tissue were measured using the Pierce BCA Protein Assay Kit (Thermo Fisher Scientific, Waltham, MA, USA).

### 2.8 Histological Analysis

Cryopreserved hemispheres were sectioned at 35-μm thickness, collected in a series of 12 into cryoprotectant solution using a freezing microtome and stored at -20°C. Immunohistochemistry was performed on one of the series with washes between each step. After rehydrating in Tris buffered saline (TBS) for 3 × 5 min, endogenous peroxidase activity was blocked by applying 1% hydrogen peroxide (H_2_O_2_) in TBS for 30 min at room temperature (RT), followed by a non-specific binding block with 10% skimmed milk powder in TBS with 2% Triton-X (TBS-X) for 2 h at RT. Sections were then incubated in primary antibody for microglia (rabbit anti-Iba1, 1:2,000, 019-19741, Alpha Laboratories, UK) diluted in 5% skimmed milk powder in TBS-X overnight at 4°C. Following three washes in TBS-X, sections were incubated in biotinylated goat anti-rabbit diluted in TBS-X (1:1,000; BA-1000; Vector Laboratories Ltd., Burlingame, USA) for 2 h at RT followed by incubation in avidin-horseradish peroxidase complex (Vectastain ABC Elite, PK-6000; Vector Laboratories) for 1 h. To confirm the specificity of the immunoreactivity, we performed the ABC incubation with no primary or secondary antibody as negative controls alongside all immunohistochemical procedures; all negative controls showed no specific staining.

Immunoreactivity was visualized by incubating sections in 0.05% diaminobenzidine and 0.01% H_2_O_2_ for up to five minutes with exact timing being determined by the depth of color of the sections; all sections were incubated at the same time and using the same timing. Sections were then rinsed in TBS, mounted on superfrost microscope slides, and left to dry overnight before dehydrating in increasing concentrations of industrial methylated spirits (IMS) followed by xylene before cover-slipping with DPX mounting medium (Sigma Aldrich).

Slides were scanned at 40× optic resolution in brightfield mode using 80% compression on export using an Olympus VS120 slide scanner (Olympus Life Science, Waltham, MA, USA). Two regions – the parietal association cortex (PtA) and the hippocampus – were identified from the MRI data as being particularly affected by LPS administration. Another two regions – the prefrontal cortex (PFC) and striatum – were identified as minimally affected by LPS administration based on the MRI data. Analysis of Iba1-positive (Iba1+) cells in these four ROIs was conducted using an adapted version (Polsek et al., 2020) of the optical fractionator method described by West et al. (West et al., 1991). First, a MATLAB script was used to place a systematic random sampling grid of 300×300 μm over manually drawn ROIs, from which jpeg images of each square of the grid were captured, together with a scale bar and a text file containing ROI area information.

Next, using a stereological tool STEPanizer (Stepanizer.com, (Tschanz et al., 2011)), a counting frame of 50 µm × 50 µm was applied to each jpeg in which Iba1+ cells were counted according to the principles of unbiased stereological estimation (Mandarim-de-Lacerda and Del Sol, 2017; West et al., 1991). The counts were converted to population estimates, ROIs were converted to volume using the Cavalieri principle, and Gundersen’s coefficient of error (CE) for the counts was calculated. CE was less than 0.1 for all animals, hence all were included in the analysis. The population estimate and ROI volume were then used to normalize the counts as cells/mm^3^ that are reported here.

Thresholding was also performed on the jpeg images created by the MATLAB script. The pre-processed images were binarized into Iba1+ cells and background using an automated custom ImageJ macro, and Euclidean distance maps were computed in which the value of each intracellular pixel denotes the shortest distance between that pixel and the cell boundary. For each rat and ROI, all these intracellular distances were summed and normalized by the total number of pixels in the images. This measure, created in-house, is a shape-weighted area fraction (SWAF) that accounts for cell size and shape. **Supplemental Figure 2** illustrates the rationale for using SWAF: it can better differentiate reactive from non-reactive microglia than standard area fraction, but it was found to correlate strongly with standard area fraction.

Mann-Whitney U tests were used to test for differences in SWAF and cellular density between groups, and the Benjamini-Hochberg procedure used to correct for multiple comparisons across stains, parameters, and ROIs. Spearman correlation coefficients between non-imaging measures (weight, locomotor activity, plasma and brain protein levels) and the Iba1+ SWAFs and median diffusion parameter values were computed for both groups pooled together and for each group separately.

### 2.9 Identification of LPS non-responders

Systemic administration of LPS does not elicit a response in approximately 10% of rats (unpublished data). Here, LPS non-responders were identified based on changes in weight and distance moved during the open field test, and plasma levels of α-2-macroglobulin. All 32 rats were classified into one of two groups by using MATLAB to perform K-means clustering on these three measures. LPS-treated rats that were clustered with the controls were considered non-responders and excluded from statistical comparisons between groups.

## 3 Results

### 3.1 Systemic LPS administration induced neuroinflammation

Intraperitoneal administration of LPS elicited an immune response characterized by sickness behavior (i.e., reduced locomotor activity and food and water consumption) and consequent weight loss. Four LPS-treated rats were identified as non-responders and excluded from the study based on the results of K-means clustering (**Figure 1a**), bringing the final group sizes to 16 LPS and 12 controls. No control rats were clustered with the LPS rats. Both systemic and neuroinflammation were confirmed in LPS responders by significantly elevated (FDR-corrected p < 0.005) plasma and brain levels of several pro-inflammatory proteins including interleukins IL1b and IL6 (**Figure 1b-g**).

**Figure 1.**
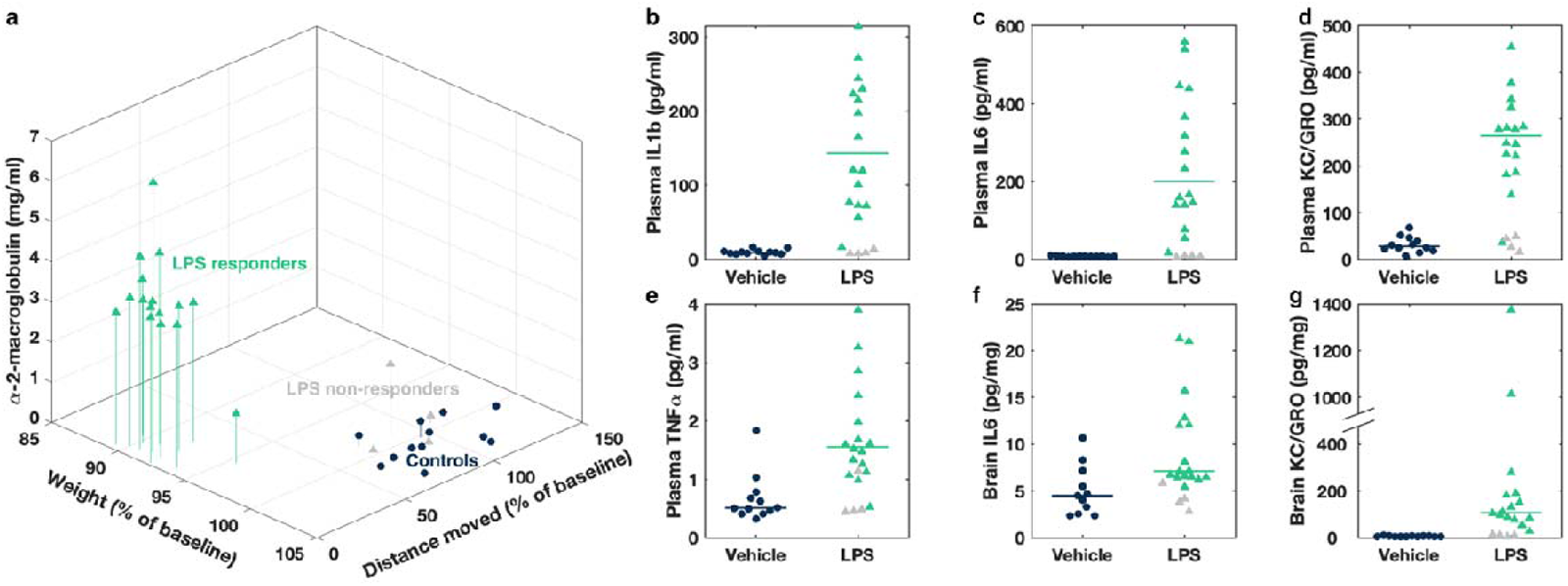
a) A 3D stem plot of weight, distance moved, and plasma levels of α-2-macroglobulin of individual rats, measured one day after i.p. injection of LPS (green triangles) or saline vehicle (blue circles). There are two distinct clusters, with the LPS non-responders (gray triangles) grouped with the control rats. Weight and distance moved are reported as percentage of baseline measurements taken before injections. b-g) Pro-inflammatory cytokine levels in the plasma (b-e) and brain (f, g). Each symbol indicates an individual rat, and horizontal lines indicate group medians. LPS non-responders are colored gray and were not included in the calculation of the LPS group medians.

### 3.2 LPS caused voxel-wise increase in water diffusivity throughout the brain

Voxel-based analysis revealed that FA was significantly decreased and MD significantly increased (FWE-corrected p < 0.05) in parts of the PtA, visual, retrosplenial, motor, and sensory cortices of LPS-treated rats compared to vehicle-treated controls (**Figure 2a, d, Supplemental Table 1**).

There was little to no effect of LPS on FAt or MDt (**Supplemental Table 1**). However, FW was significantly greater in areas of the cortex in LPS-treated rats, displaying a similar spatial pattern to MD (**Figure 2g**, **Supplemental Table 1**).

While f_iso_ ostensibly measures the same thing as FW and correlated very strongly with FW (median Spearman rho ρ = 0.93, **Supplemental Figure 4**), it was not significantly affected by LPS administration (**Supplemental Table 1**). In comparison, f_hin_ was significantly increased in larger areas of the cortex as well as parts of the corpus callosum (**Figure 2h**, **Supplemental Table 1**). f_*res*_ was the most extensively affected, being significantly decreased in the LPS group in even greater proportions of these structures than f_hin_, in addition to various subcortical regions including the hippocampus, thalamus, and basal forebrain. (**Figure 2i**, **Supplemental Table 1**). ODI was not significantly different between groups (**Supplemental Table 1**).

To better understand the effects of LPS on diffusivity in the brain, DTI modeling was performed on each of the three diffusion shells separately. MD and MD500 correlated most strongly with FW (median ρ = 0.86 and 0.73, respectively) while MD1000 and MD1500 correlated most strongly with f_*res*_ (median ρ = -0.72 and -0.78, 17 respectively) (**Supplemental Figure 4**). MD500 was significantly increased in various cortical areas beyond where FW was increased, as well as in parts of the corpus callosum and dorsal hippocampus (**Figure 2e**, **Supplemental Table 1**). MD1500 was significantly elevated in many of the same areas in which f_*res*_ was decreased (**Figure 2f**, **Supplemental Table 1**). Interestingly, the greatest decrease in fractional anisotropy was seen when using the b500 shell only (**Figure 2b**, **Supplemental Table 1**). There was no significant between-group difference in FA1000 or MD1000 (**Supplemental Table 1**).

**Figure 2.**
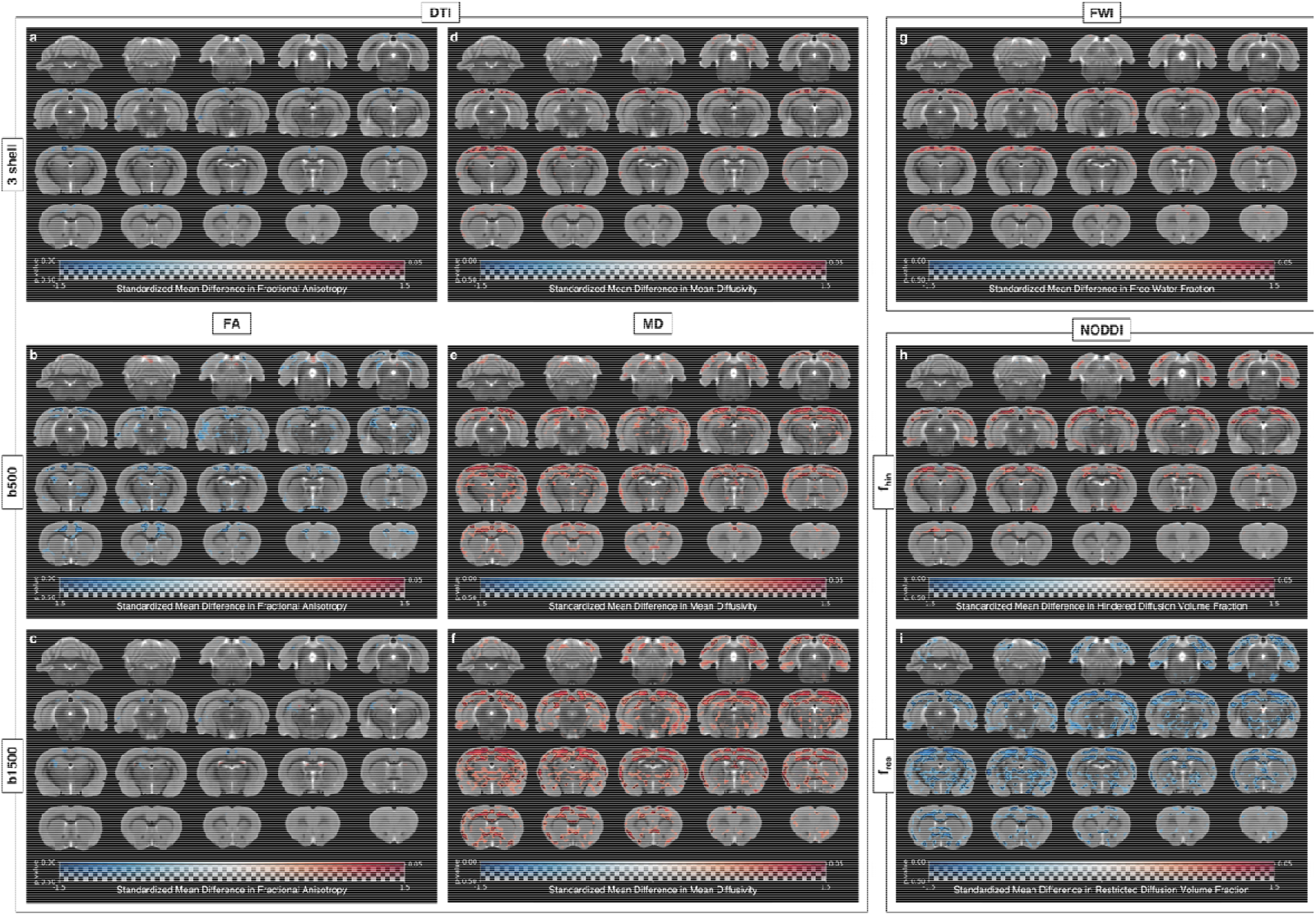
Dual-coded maps showing voxel-wise differences in dMRI parameters between LPS and vehicle groups, overlaid on coronal slices of a T2-weighted rat brain template (Valdes-Hernandez). The standardized mean difference between groups is coded by overlay color (red indicates the parameter is larger in the LPS group), and the statistical significance is coded by overlay transparency (greater opacity indicates greater statistical significance). The overlay is completely transparent in regions in which the family-wise-error (FWE)-corrected p > 0.5; black lines circumscribe regions in which FWE-corrected p < 0.05. a-c) Fractional anisotropy calculated from the simple DTI model using a) all three shells, b) just the b500 shell, and c) just the b1500 shell. d-f) Mean diffusivity calculated from the simple DTI model using a) all three shells, b) just the b500 shell, and c) just the b1500 shell. d) Free water volume fraction calculated from the FWI model. e) Hindered diffusion volume fraction (f_*hin*_) and f) restricted diffusion volume fraction (f_*res*_) calculated from the NODDI model.

### 3.3 LPS-induced changes in diffusion were most significant in the parietal association cortex

The analysis of 59 atlas-based anatomical ROIs confirmed that FW, f_*res*_, MD500, and MD1500 were most significantly affected in the PtA by LPS administration (**Figure 3, left column**). The visual and retrosplenial cortices are two other ROIs in which diffusion parameters were highly affected (**Figure 3, middle and right columns**). **Table 1** lists the 15 ROIs with the most significant differences in FW, f_*res*_, MD500, and MD1500 between LPS and control groups. Median f_*res*_ was significantly lower in 15 ROIs (FDR-corrected p < 0.05), median MD1500 significantly greater in 12 ROIs, and median MD500 significantly greater in seven ROIs. FW was not significantly different in any ROI. Most of the most affected regions were in the gray matter, but some white matter regions were also significantly affected: corpus callosum and descending corticofugal pathways (f_*res*_ and MD1500), stria terminalis (f_*res*_), and ventral hippocampal commissure (MD500 and MD1500).

**Figure 3.**
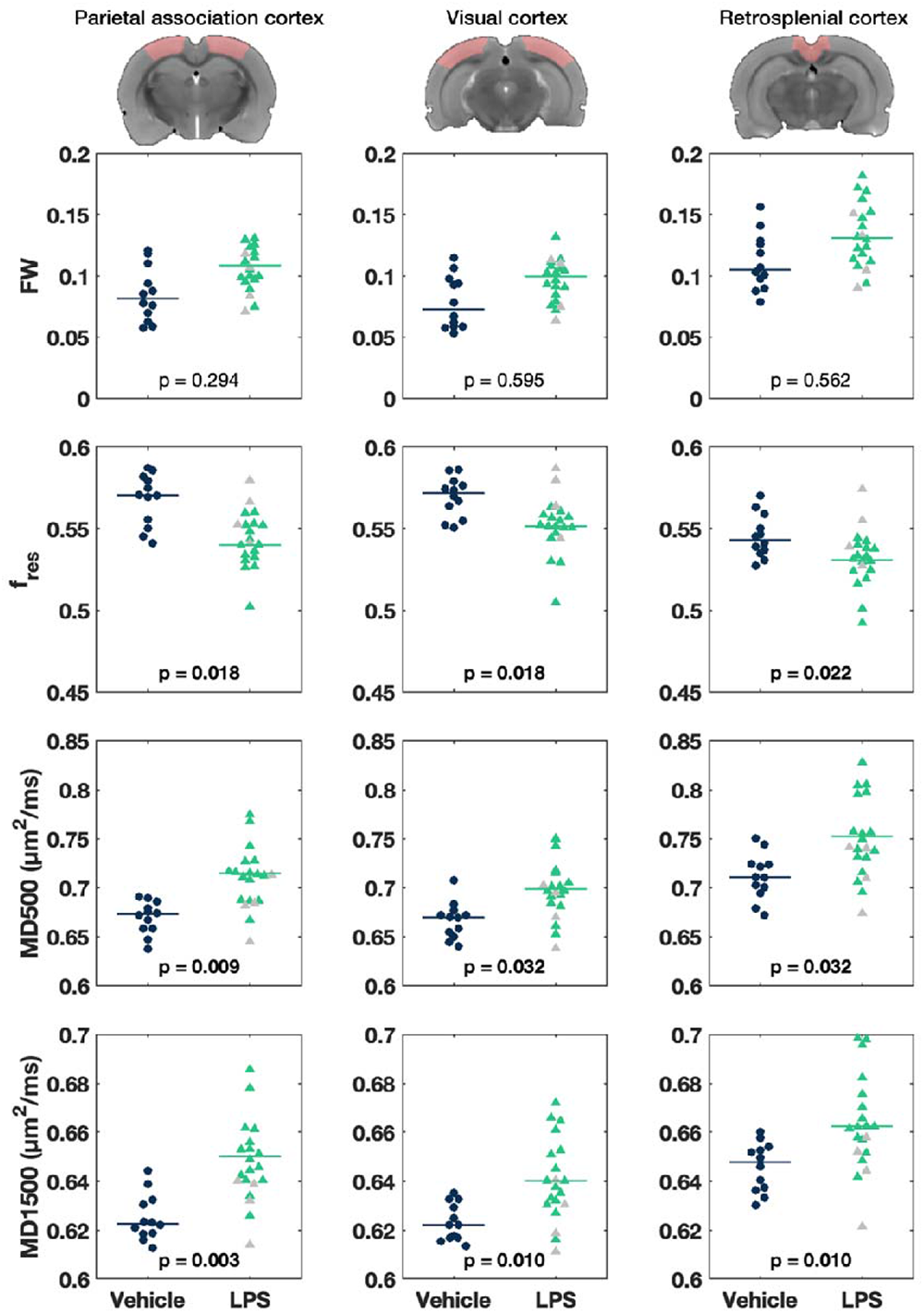
Median dMRI parameter values in three select atlas regions-of-interest: parietal association cortex (left column), visual cortex (middle column), and retrosplenial cortex (right column). FW: free water volume fraction (FWI); fres: restricted diffusion volume fraction (NODDI); MD500: b500 mean diffusivity, and MD1500: b1500 mean diffusivity (DTI). Each symbol indicates an individual rat, and horizontal lines indicate group medians. P-values are from two-tailed Mann-Whitney U tests and FDR-corrected. Gray triangles: LPS non-responders, excluded from statistical tests and calculation of LPS group medians.

**Table 1:**
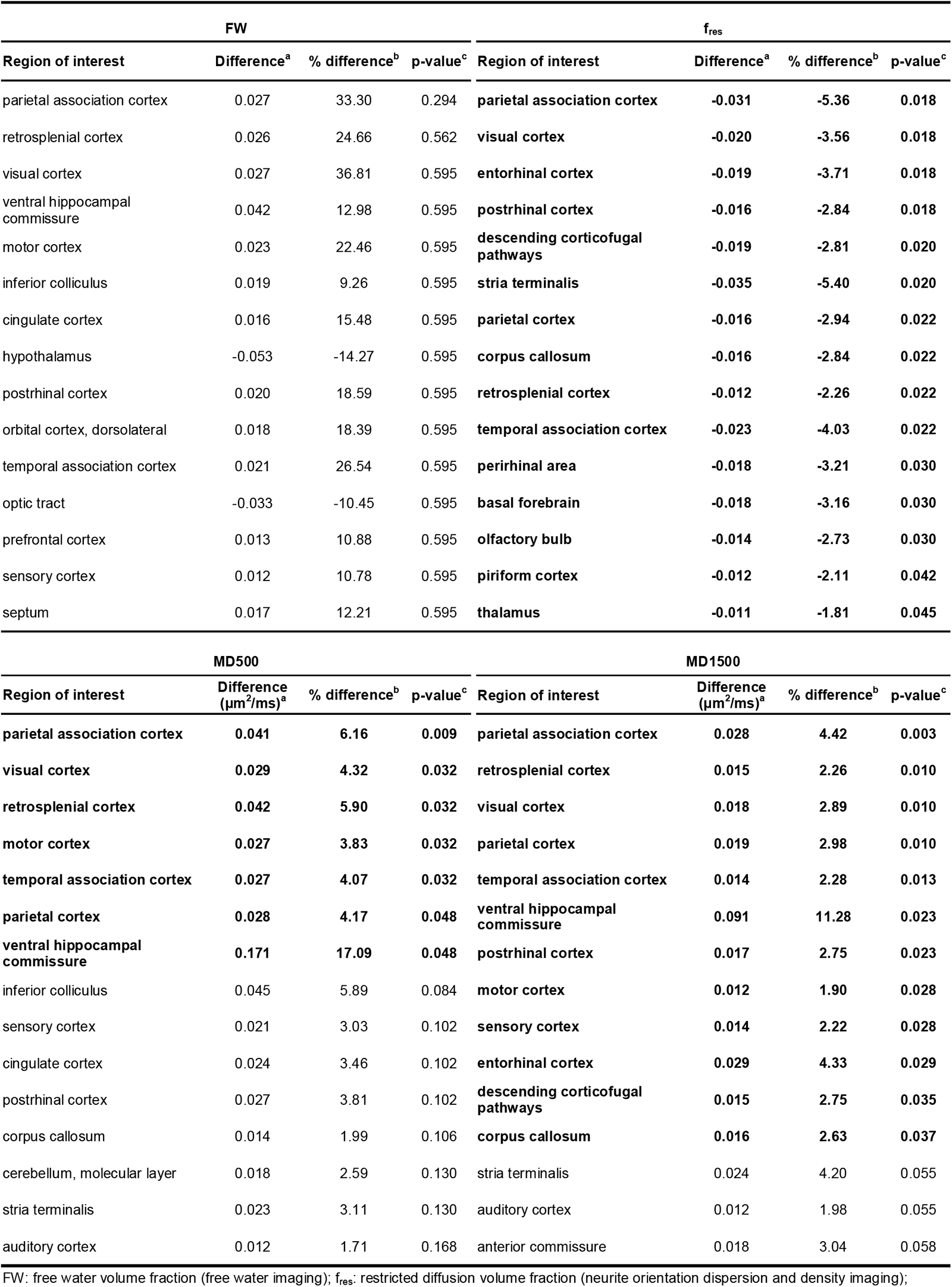
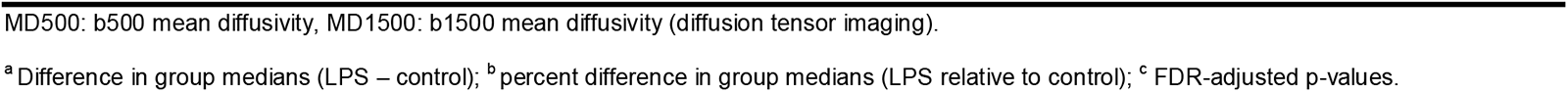
ROIs with the most significant differences in dMRI parameters between LPS-treated and control rats, ordered by increasing p-value.

### 3.4 Increased diffusivity correlates with changes in microglial morphology

Iba1 immunohistochemistry revealed microglial activation in the PtA, hippocampus, striatum, and PFC of LPS-treated rat brains (**Figure 4a**). This was quantified by a significantly increased SWAF of Iba1+ cells (p < 0.05; indicative of larger, more ameboid shapes) (**Figure 4b**). There were also significantly more Iba1+ cells in the hippocampus of LPS rats compared to controls (p = 0.0038, **Figure 4c**, second row) but not in the other three ROIs (p > 0.4). GFAP+ SWAF, GFAP+ cell density, and Nissl+ cell density were measured in the PtA and hippocampus but were not significantly different between LPS and vehicle groups in either ROI (**Supplemental Figure 5**).

**Figure 4.**
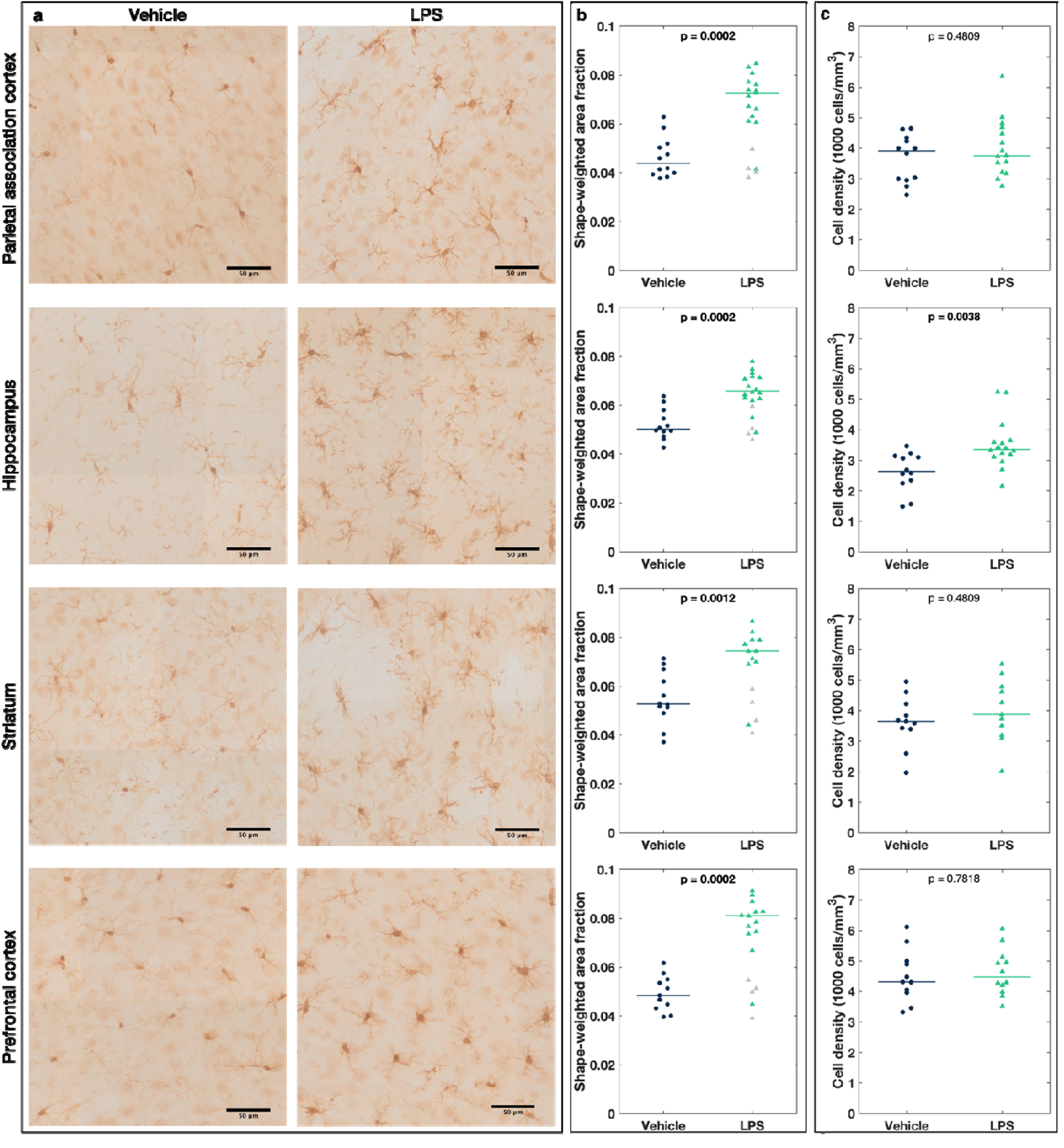
a) Representative 40× images of Iba1-stained brain sections from the parietal association cortex (first row), hippocampus (second row), striatum (third row), and prefrontal cortex (fourth row) of a vehicle-treated control (left column) and LPS-treated rat (right column). b) Shape-weighted area fraction of Iba1+ cells. c) Stereologically estimated Iba1+ cell density in the same four regions of individual rats. Horizontal lines indicate group medians. P-values are from two-tailed Mann-Whitney U tests and FDR-corrected. Gray triangles: LPS non-responders, excluded from statistical tests and calculation of LPS group medians.

**Table 2** shows the Spearman correlation coefficients between the histologically measured Iba1+ SWAFs and the median values of the dMRI parameters FW, f_*res*_, MD500, and MD1500 in the PtA, hippocampus, striatum, and PFC. Correlation coefficients were computed for each group separately as well as for all rats combined. No significant within-group correlations were found except with MD1500 in the PtA in LPS-treated rats. When both groups were combined, Iba1+ SWAF correlated significantly with all four diffusion parameters in the PtA and with f_*res*_ and MD1500 in the hippocampus. Iba1+ SWAF correlated inversely with f*_res_* and directly with the other diffusion parameters. There was no correlation between Iba1+ SWAF and any diffusion parameter in the striatum or PFC. Scatter plots of median FW, f_*res*_, MD500, and MD1500 vs. Iba1+ SWAF are shown in **Figure 5**.

**Table 2:**
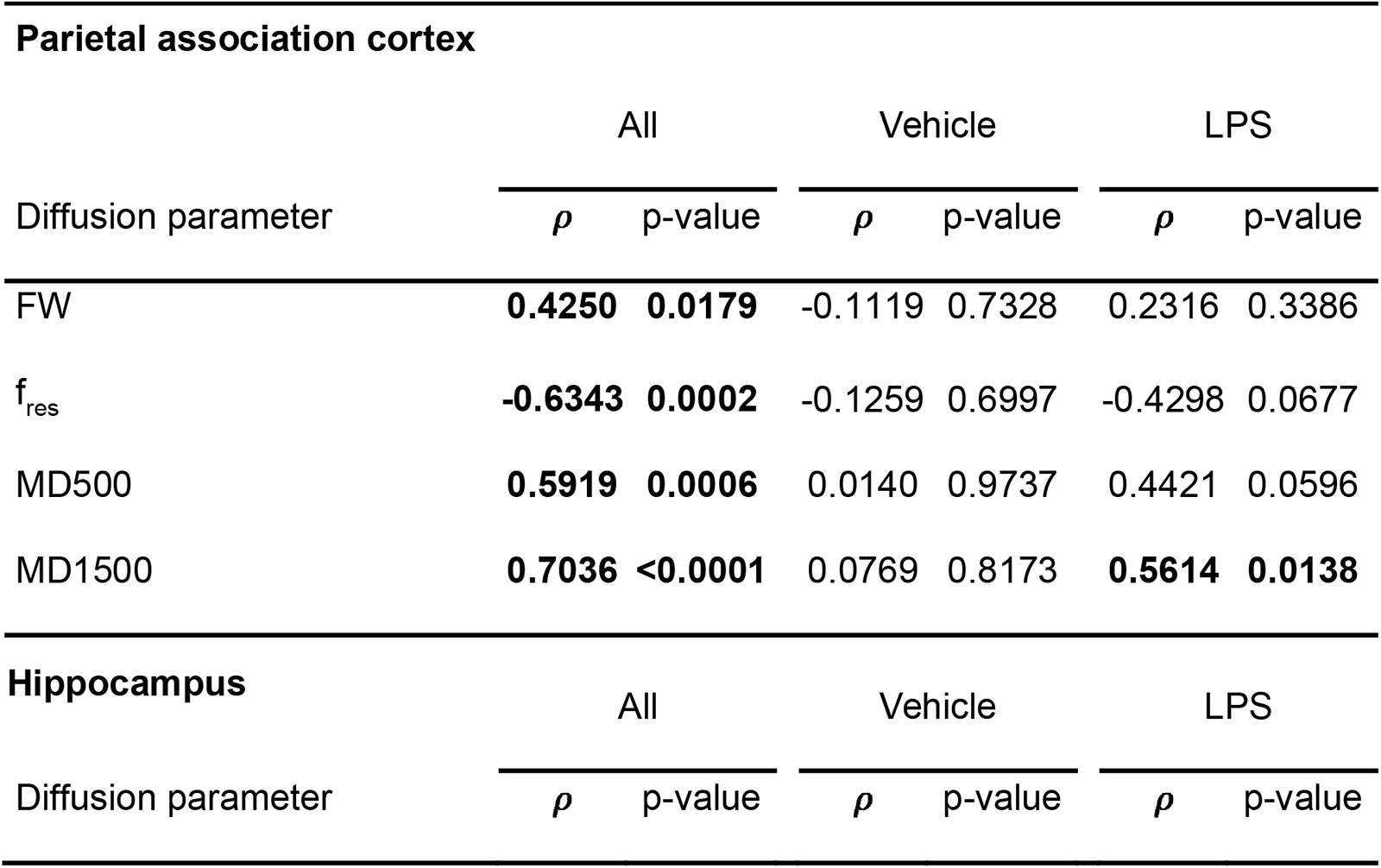
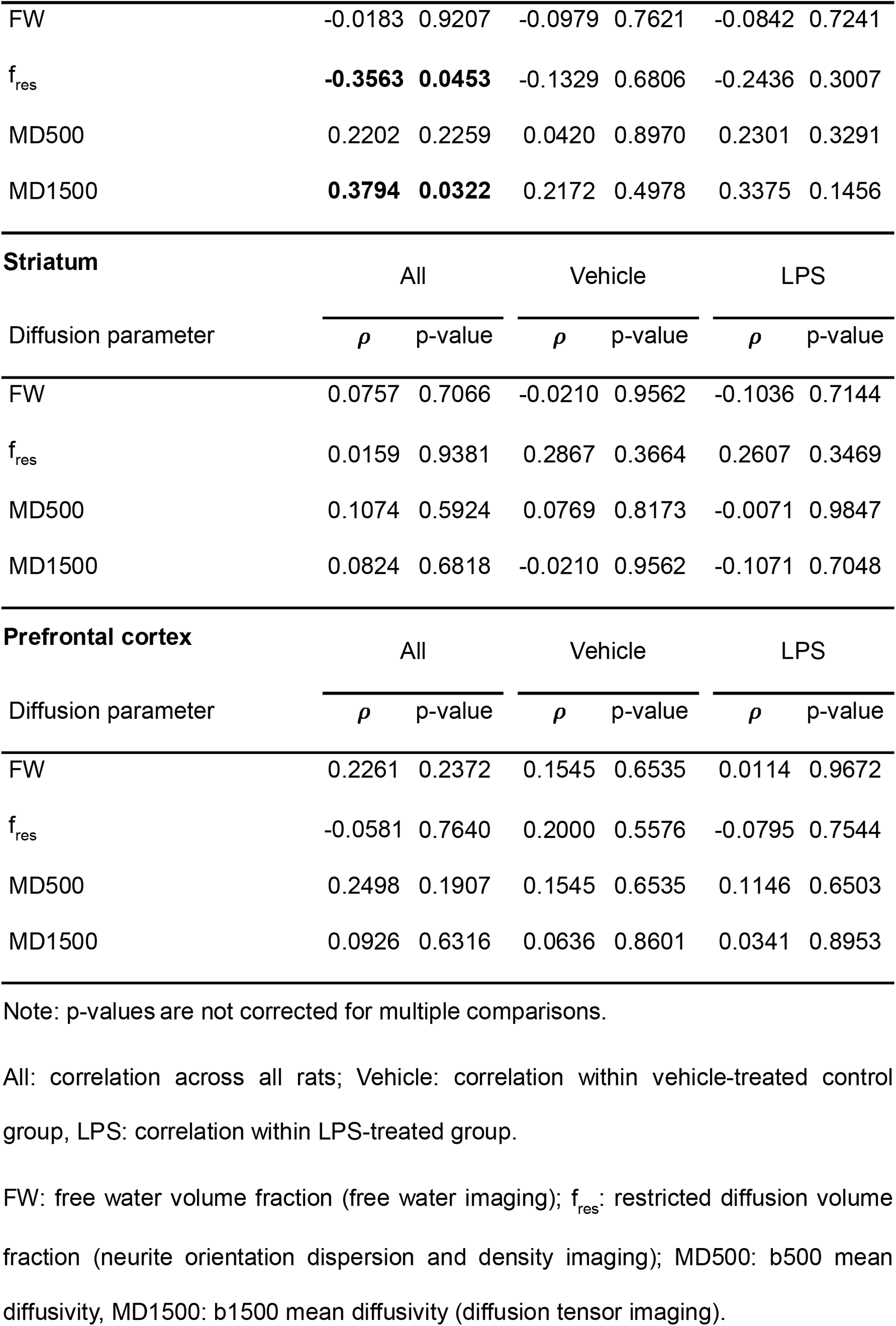
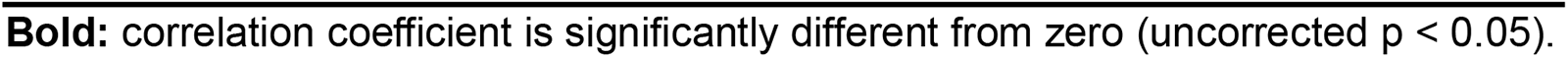
Spearman correlations between dMRI parameters and Iba1+ SWAF in select ROIs

**Figure 5.**
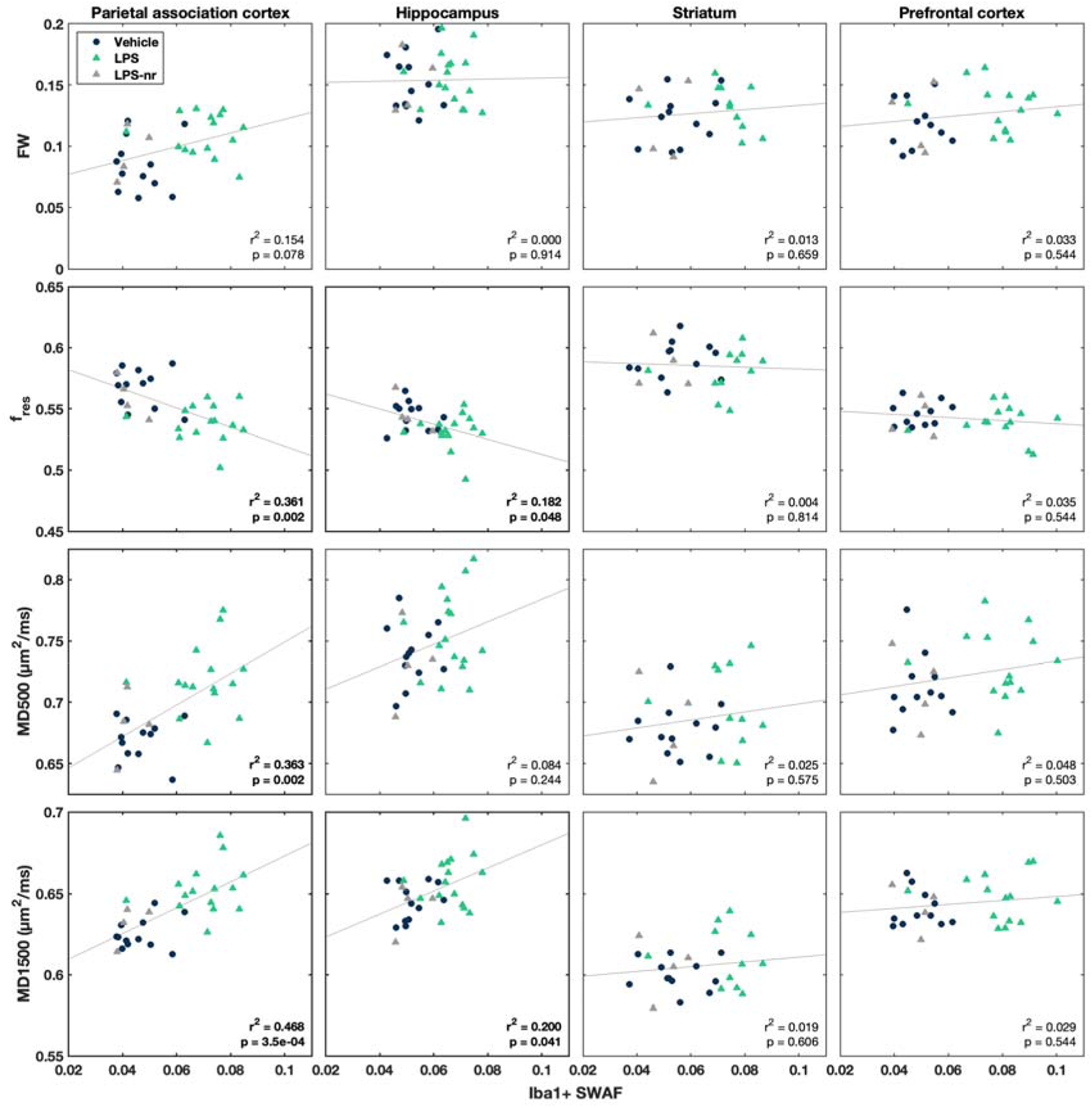
Median dMRI parameters vs. histologically measured Iba1+ shape-weighted area fraction (SWAF) in the parietal association cortex, hippocampus, striatum, and prefrontal cortex. Each symbol represents an individual rat. Blue circles represent control rats, green triangles LPS rats, and gray triangles LPS non-responders. Linear regression lines (all rats pooled) are plotted in gray, and the R-squared statistic and uncorrected p-value of each regression model displayed in each plot. FW: free water volume fraction (free water imaging); fres: restricted diffusion volume fraction (neurite orientation dispersion and density imaging); MD500: b500 mean diffusivity, MD1500: b1500 mean diffusivity (diffusion tensor imaging).

### 3.5 Diffusivity and microglial morphology correlate with behavior and cytokine levels

Scatter plots of Iba1+ SWAF and MD1500 in the PtA vs. non-imaging measures (weight, locomotor activity, plasma and brain protein levels) are shown in **Supplemental Figure 6**. The corresponding Spearman correlation coefficients are reported in **Supplemental Table 2**. With all rats pooled, SWAF and MD1500 correlated significantly with all nine non-imaging measures. Within the LPS-treated group but without correcting for multiple comparisons, SWAF correlated significantly with weight loss and plasma levels of α-2-macroglobulin and IL1b, while MD1500 correlated significantly with locomotor activity (change in distance moved) and plasma levels of IL1b and IL6 (uncorrected p < 0.05).

## 4 Discussion

### 4.1 LPS increased brain diffusivity

In this study, *in vivo* diffusion MRI was used to detect acute neuroinflammation induced by systemic LPS administration in rats. Neuroinflammation was confirmed by increased pro-inflammatory cytokines and reactive microglia in the brain. Three diffusion modelling techniques – DTI, FWI, and NODDI – were employed to quantify the inflammation-induced changes on the brain tissue microstructure. All three models were sensitive to LPS-induced changes, together indicating an increase in water diffusivity because of increased extracellular and/or microglial cell volume.

The apparent spatial extent and magnitude of the LPS response differed between diffusion metrics. The greater increases in MD500 and MD1500 compared to MD and MD1000 point to a larger effect of LPS-induced inflammation on the fast- and slow-diffusing water populations than on the intermediate population, and perhaps to two distinct neuroinflammatory phenomena. The increase in MD500 suggests increased extracellular water while the increase in MD1500 may be the result of microglial enlargement. Although it has been suggested that fast and slow diffusion correspond to extra- and intracellular compartments, respectively, these two populations do not map to distinct tissue structures (Le Bihan, 2013), and intracellular diffusion has been shown to have both fast and slow components (Sehy et al., 2002). Further in-depth investigations are required to explore how dMRI can characterize the complex, multifaceted processes involved in neuroinflammation.

### 4.2 Regional variation in diffusion changes

Overall, the greatest effects were detected in primarily cortical gray matter regions, but with some involvement of subcortical and white matter areas. The most affected was the parietal association cortex in which parameters from all three modalities were correlated with microglial morphology and density (SWAF) – a proxy for microglia reactivity – measured using corroborative histology. Interestingly, we detected reactive microglia in several other brain regions – notably the hippocampus, striatum, and prefrontal cortex – where there were smaller or no detected changes in diffusion, and correlations were largely absent except in the hippocampus where f*_res_* and MD1500 showed a significant but weak correlation with SWAF (**Table 2**, **Figure 5**).

This regional discrepancy between histology and MRI is an interesting question that requires further investigation. A potential contributing factor may be the inhomogeneous sensitivity profile of the surface receiver coil – the lower signal-to-noise ratio farther from the coil may have obfuscated subtle changes in subcortical brain regions. Furthermore, it has recently been shown in human DTI data that the reproducibility and reliability of diffusion metrics vary across different brain structures (Luque Laguna et al., 2020). It is also possible that intrinsic regional microstructural differences in cells such as microglia (Tan et al., 2020), or gray matter in general (Herculano-Houzel, 2012), affect diffusion model fitting and diffusion metrics in a way that results in the differential sensitivity to detect changes in specific brain areas.

For example, the hippocampus exhibited robust microglial response and recruitment, but only focal diffusion changes. Previous work has indicated that some diffusion metrics in the rodent hippocampus diverge from the established and expected parameters: Stolp et al. showed a positive correlation of MD with both cellular and axonal density (Stolp et al., 2018), which is counterintuitive as MD is generally seen to be low in areas with high axonal density. Indeed, a recent study by Radhakrishnan et al. highlighted these important regional variations by demonstrating multiple different patterns of correlations between cell types and diffusion metrics across the mouse brain (Radhakrishnan et al., 2022). Interestingly, while microglia were not significantly correlated with any diffusion metrics in many gray matter areas, they were highly correlated in the somatosensory cortex, an area that showed some of the greatest LPS-induced diffusion changes in our study. It is out the scope of this paper to investigate the interactions between regional brain architecture and diffusivity metrics, but our results point toward the need for greater understanding of such parameters before MRI can deliver reliable biomarkers of neuroinflammation.

### 4.3 The effect of microglia on diffusion

Recent work by Yi et al. demonstrated that *ex vivo* NODDI imaging is sensitive to microglia depletion in mouse brains following CSF1R inhibition, and to subsequent repopulation after withdrawal of the inhibitor (Yi et al., 2019). Interestingly, they found that ODI was significantly decreased in the dentate gyrus compared to control animals, which they attributed to decreased hindered diffusion resulting from decreased glial density; but no significant differences in FA or MD were found, while the other NODDI parameters were not reported. In contrast, our study, conducted in live animals, found significant differences in FA, MD, f_*hin*_, and f_*res*_, but not in ODI. This discrepancy may be explained by the dramatic difference in the magnitude of the responses to CSF1R inhibitor vs LPS – the former resulted in almost complete depletion of microglia while the latter resulted in morphological changes but relatively little effect on microglial density (**Figure 4**). ODI reflects the degree of dispersion in the orientation of neurites (and presumably glial processes as well). Decreased ODI is consistent with loss of highly orientationally disperse microglial processes through microglial depletion or process retraction by reactive microglia in chronic inflammation. However, our Iba1 staining revealed highly ramified microglia 24 h after LPS administration (**Figure 4**), and thus ODI was not expected to change during acute LPS-mediated neuroinflammation. There was also little effect on neurons and astrocytes as confirmed by Nissl and GFAP staining, respectively (**Supplemental Figure 5**).

Given the histological results and the lack of significant change in f_iso_, the decreased f_*res*_ we observed in LPS-treated rats is likely due to an increase in the volume of the hindered diffusion compartment, which ostensibly corresponds to the extra-neurite space including soma, glial cells, and extracellular space. This is consistent with the transformation of microglia into larger, more amoeboid shapes upon activation (Davis et al., 1994). The colocalized increase in MD1500 suggests that diffusivity is increased within swollen, reactive microglia due to increased intracellular distances between cell membranes. This is supported by the correlations found between f_*res*_, MD1500 and the histologically derived Iba1+ SWAF.

However, changes in microglial morphology upon activation are likely not the only driver of the changes in diffusion measured in LPS-treated rats. The lack of intra-group correlations between dMRI parameters and Iba1+ SWAF suggests that the observed changes in these diffusion parameters were influenced by another effect. Although difficult to verify, a possible explanation is mild vasogenic edema due to the neuroinflammatory response mediated by reactive microglia and other immune cells.

Garcia-Hernandez et al. recently demonstrated that *in vivo* diffusion MRI is sensitive to changes in microglial morphology in rats following an intracerebral LPS challenge (Garcia-Hernandez et al., 2022). They present a model of gray matter diffusion comprised of four compartments: one cylindrically restricted diffusion compartment representing cellular processes; one small and one large spherically restricted diffusion compartments representing microglial and astrocytic somas, respectively; and an extracellular compartment modeled as a tensor aligned to the main cylinder orientation of the first compartment. At eight and 24 hours after intracerebral administration of LPS into the hippocampus of one hemisphere, the stick fraction (similar to NODDI f_*res*_) was decreased, and the small sphere radius (ostensibly the microglial soma size) was increased in the region that received LPS compared to the contralateral hippocampus that received saline. These findings are consistent with the decreased f_*res*_ and increased f_*hin*_ in LPS-treated rats found in the current study. The stick fraction was shown to correlate strongly with microglial process density measured from Iba1 histology, and LPS-induced changes in stick fraction and small sphere radius were reduced after microglial depletion by CSF1R inhibition, demonstrating the specificity of these diffusion parameters to microglial morphology.

Although these results are very promising, the study is somewhat limited by the use of non-physiological CSF1R inhibitor and LPS models. It remains to be seen if this method is sensitive to less invasive and therefore more clinically translational responses. Moreover, the complex modelling requires the acquisition of images with multiple diffusion times as well as multiple shells, which increases the minimum scan time and limits routine implementation for researchers who wish to study neuroinflammation or its treatments in various experimental settings.

### 4.4 Choice of diffusion model

Diffusion MRI and the DTI model remain important and powerful tools to identify subtle neuroinflammatory abnormalities (O’Donnell and Pasternak, 2015). Indeed, the recent study by Garcia-Hernandez et al. and this current study demonstrate that simple MD can be as sensitive to neuroinflammation as more complex diffusion models (Garcia-Hernandez et al., 2022). However, FA and MD are also sensitive to many other structural changes, such as demyelination, membrane permeability, changes in fiber organization, and partial volume effects (Alexander et al., 2007; Jones et al., 2013). More complex models may be a solution to DTI’s lack of specificity, but model fitting becomes noisier and less robust as more free parameters are added – more and higher quality data is required to obtain reliable fits. Thus, the pursuit of sensitive and specific biomarkers of neuroinflammation is an ongoing challenge.

### 4.5 Study limitations

Our study has several limitations. First, the b-values used were lower than what is recommended for NODDI (Zhang et al., 2012). Our rationale was two-fold: 1) the need to better characterize fast diffusion components where we expected to see greater effects of inflammation, and 2) because pilot data at high b-value was considered of poor quality due to motion artifacts even with respiratory gating. Thus, it was decided to focus only on lower b-values considering that we did not expect to see changes in highly restricted compartments.

The LPS challenge was administered, and the behavioral tests performed during the lights-on phase (i.e., the animals’ inactive/sleep) period. While this is common in animal research, it bears mentioning that this can impact the translatability of preclinical studies.

Although here we used a relatively non-invasive systemic LPS challenge to induce neuroinflammation (e.g., compared to intracerebral LPS injection), this still elicits a large and acute inflammatory response. To make these results more clinically translatable, it would be important to verify and extend to other more chronic neuroinflammatory conditions such as, e.g., those related to stress or maternal immune activation.

Moreover, it is important to corroborate the results by histological methods, but there is an inherent difficulty in comparing diffusion imaging with post-mortem techniques. This is because the former relies on measuring the behavior of water molecules, but histological fixation removes and artificially rearranges tissue water and extra/intracellular spaces (Schilling et al., 2022). While this and previous studies have demonstrated relationships between cellular architecture and dMRI measurements, using histology to corroborate the effects of extracellular changes (often hypothesized in neuroinflammatory conditions) is an unmet challenge.

### 4.6 Conclusions and future work

Using a combination of three types of diffusion MRI, and a reproducible and translational scan protocol, we show for the first time the changes in rat brain diffusivity *in vivo* in a rodent model of acute low grade neuroinflammation that is relevant to sickness behavior and depression. Lack of specificity is a limitation of MRI in general, but here we demonstrate that diffusion MRI is sensitive to microstructural changes that accompany neuroinflammation. Future work should include implementing more sophisticated diffusion models to improve specificity and leveraging the sensitivity of current methods to evaluate treatment response.

## Supporting information

Appendix - NIMA Members

## Data availability

Raw data is available on OpenNeuro (https://doi.org/10.18112/openneuro.ds004305.v1.0.1).

## References

Agosta, F., Pievani, M., Sala, S., Geroldi, C., Galluzzi, S., Frisoni, G.B., Filippi, M., 2011. White matter damage in Alzheimer disease and its relationship to gray matter atrophy. Radiology 258, 853–863.

Albrecht, D.S., Granziera, C., Hooker, J.M., Loggia, M.L., 2016. In Vivo Imaging of Human Neuroinflammation. ACS Chem Neurosci 7, 470–483.

Alexander, A.L., Lee, J.E., Lazar, M., Field, A.S., 2007. Diffusion tensor imaging of the brain. Neurotherapeutics 4, 316–329.

Althubaity, N., Schubert, J., Martins, D., Yousaf, T., Nettis, M.A., Mondelli, V., Pariante, C., Harrison, N.A., Bullmore, E.T., Dima, D., Turkheimer, F.E., Veronese, M., 2022. Choroid plexus enlargement is associated with neuroinflammation and reduction of blood brain barrier permeability in depression. Neuroimage Clin 33, 102926.

Andersson, J.L., Skare, S., Ashburner, J., 2003. How to correct susceptibility distortions in spin-echo echo-planar images: application to diffusion tensor imaging. Neuroimage 20, 870–888.

Andersson, J.L.R., Sotiropoulos, S.N., 2016. An integrated approach to correction for off-resonance effects and subject movement in diffusion MR imaging. Neuroimage 125, 1063–1078.

Avants, B.B., Epstein, C.L., Grossman, M., Gee, J.C., 2008. Symmetric diffeomorphic image registration with cross-correlation: evaluating automated labeling of elderly and neurodegenerative brain. Med Image Anal 12, 26–41.

Avants, B.B., Tustison, N.J., Song, G., Cook, P.A., Klein, A., Gee, J.C., 2011. A reproducible evaluation of ANTs similarity metric performance in brain image registration. Neuroimage 54, 2033–2044.

Batista, C.R.A., Gomes, G.F., Candelario-Jalil, E., Fiebich, B.L., de Oliveira, A.C.P., 2019. Lipopolysaccharide-Induced Neuroinflammation as a Bridge to Understand Neurodegeneration. Int J Mol Sci 20.

Biesmans, S., Meert, T.F., Bouwknecht, J.A., Acton, P.D., Davoodi, N., De Haes, P., Kuijlaars, J., Langlois, X., Matthews, L.J., Ver Donck, L., Hellings, N., Nuydens, R., 2013. Systemic immune activation leads to neuroinflammation and sickness behavior in mice. Mediators Inflamm 2013, 271359.

Bowman, G.L., Quinn, J.F., 2008. Alzheimer’s disease and the Blood-Brain Barrier: Past, Present and Future. Aging health 4, 47–55.

Chang, L., Munsaka, S.M., Kraft-Terry, S., Ernst, T., 2013. Magnetic resonance spectroscopy to assess neuroinflammation and neuropathic pain. J Neuroimmune Pharmacol 8, 576–593.

Collste, K., Forsberg, A., Varrone, A., Amini, N., Aeinehband, S., Yakushev, I., Halldin, C., Farde, L., Cervenka, S., 2016. Test-retest reproducibility of [(11)C]PBR28 binding to TSPO in healthy control subjects. Eur J Nucl Med Mol Imaging 43, 173–183.

Dantzer, R., O’Connor, J.C., Freund, G.G., Johnson, R.W., Kelley, K.W., 2008. From inflammation to sickness and depression: when the immune system subjugates the brain. Nat Rev Neurosci 9, 46–56.

Davis, E.J., Foster, T.D., Thomas, W.E., 1994. Cellular forms and functions of brain microglia. Brain Res Bull 34, 73–78.

Dhaya, I., Griton, M., Raffard, G., Amri, M., Hiba, B., Konsman, J.P., 2018. Bacterial lipopolysaccharide-induced systemic inflammation alters perfusion of white matter-rich regions without altering flow in brain-irrigating arteries: Relationship to blood-brain barrier breakdown? J Neuroimmunol 314, 67–80.

Dickie, B.R., Vandesquille, M., Ulloa, J., Boutin, H., Parkes, L.M., Parker, G.J.M., 2019. Water-exchange MRI detects subtle blood-brain barrier breakdown in Alzheimer’s disease rats. Neuroimage 184, 349–358.

Gaitan, M.I., Shea, C.D., Evangelou, I.E., Stone, R.D., Fenton, K.M., Bielekova, B., Massacesi, L., Reich, D.S., 2011. Evolution of the blood-brain barrier in newly forming multiple sclerosis lesions. Ann Neurol 70, 22–29.

Garcia-Hernandez, R., Cerdan Cerda, A., Trouve Carpena, A., Drakesmith, M., Koller, K., Jones, D.K., Canals, S., De Santis, S., 2022. Mapping microglia and astrocyte activation in vivo using diffusion MRI. Sci Adv 8, eabq2923.

Griton, M., Dhaya, I., Nicolas, R., Raffard, G., Periot, O., Hiba, B., Konsman, J.P., 2020. Experimental sepsis-associated encephalopathy is accompanied by altered cerebral blood perfusion and water diffusion and related to changes in cyclooxygenase-2 expression and glial cell morphology but not to blood-brain barrier breakdown. Brain Behav Immun 83, 200–213.

Guilarte, T.R., Rodichkin, A.N., McGlothan, J.L., Acanda De La Rocha, A.M., Azzam, D.J., 2022. Imaging neuroinflammation with TSPO: A new perspective on the cellular sources and subcellular localization. Pharmacol Ther 234, 108048.

Herculano-Houzel, S., 2012. The remarkable, yet not extraordinary, human brain as a scaled-up primate brain and its associated cost. Proc Natl Acad Sci U S A 109 Suppl 1, 10661–10668.

Heye, A.K., Culling, R.D., Valdes Hernandez Mdel, C., Thrippleton, M.J., Wardlaw, J.M., 2014. Assessment of blood-brain barrier disruption using dynamic contrast-enhanced MRI. A systematic review. Neuroimage Clin 6, 262–274.

Hoogland, I.C., Houbolt, C., van Westerloo, D.J., van Gool, W.A., van de Beek, D., 2015. Systemic inflammation and microglial activation: systematic review of animal experiments. J Neuroinflammation 12, 114.

Ilanges, A., Shiao, R., Shaked, J., Luo, J.D., Yu, X., Friedman, J.M., 2022. Brainstem ADCYAP1(+) neurons control multiple aspects of sickness behaviour. Nature 609, 761–771.

Inglese, M., Bester, M., 2010. Diffusion imaging in multiple sclerosis: research and clinical implications. NMR Biomed 23, 865–872.

Jones, D.K., Knosche, T.R., Turner, R., 2013. White matter integrity, fiber count, and other fallacies: the do’s and don’ts of diffusion MRI. Neuroimage 73, 239–254.

Karlsgodt, K.H., 2019. Using Advanced Diffusion Metrics to Probe White Matter Microstructure in Individuals at Clinical High Risk for Psychosis. Am J Psychiatry 176, 777–779.

Klawiter, E.C., Schmidt, R.E., Trinkaus, K., Liang, H.F., Budde, M.D., Naismith, R.T., Song, S.K., Cross, A.H., Benzinger, T.L., 2011. Radial diffusivity predicts demyelination in ex vivo multiple sclerosis spinal cords. Neuroimage 55, 1454–1460.

Konsman, J.P., Parnet, P., Dantzer, R., 2002. Cytokine-induced sickness behaviour: mechanisms and implications. Trends Neurosci 25, 154–159.

Kwon, H.S., Koh, S.H., 2020. Neuroinflammation in neurodegenerative disorders: the roles of microglia and astrocytes. Transl Neurodegener 9, 42.

Le Bihan, D., 2013. Apparent diffusion coefficient and beyond: what diffusion MR imaging can tell us about tissue structure. Radiology 268, 318–322.

Luque Laguna, P.A., Combes, A.J.E., Streffer, J., Einstein, S., Timmers, M., Williams, S.C.R., Dell’Acqua, F., 2020. Reproducibility, reliability and variability of FA and MD in the older healthy population: A test-retest multiparametric analysis. Neuroimage Clin 26, 102168.

Mandarim-de-Lacerda, C.A., Del Sol, M., 2017. Tips for Studies with Quantitative Morphology (Morphometry and Stereology). International Journal of Morphology 35, 1482–1494.

Miller, A.H., Haroon, E., Felger, J.C., 2017. The Immunology of Behavior-Exploring the Role of the Immune System in Brain Health and Illness. Neuropsychopharmacology 42, 1–4.

Mondelli, V., Vernon, A.C., Turkheimer, F., Dazzan, P., Pariante, C.M., 2017. Brain microglia in psychiatric disorders. Lancet Psychiatry 4, 563–572.

Nazeri, A., Schifani, C., Anderson, J.A.E., Ameis, S.H., Voineskos, A.N., 2020. In Vivo Imaging of Gray Matter Microstructure in Major Psychiatric Disorders: Opportunities for Clinical Translation. Biol Psychiatry Cogn Neurosci Neuroimaging 5, 855–864.

Nettis, M.A., Pariante, C.M., 2020. Is there neuroinflammation in depression? Understanding the link between the brain and the peripheral immune system in depression. Int Rev Neurobiol 152, 23–40.

Notter, T., Schalbetter, S.M., Clifton, N.E., Mattei, D., Richetto, J., Thomas, K., Meyer, U., Hall, J., 2021. Neuronal activity increases translocator protein (TSPO) levels. Mol Psychiatry 26, 2025–2037.

Nutma, E., Fancy, N., Weinert, M., Marzin, M.C., Tsartsalis, S., Muirhead, R.C.J., Falk, I., de Bruin, J., Hollaus, D., Pieterman, R., Anink, J., Story, D., Chandran, S., Tang, J., Trolese, M.C., Saito, T., Saido, T.C., Wiltshire, K., Beltran-Lobo, P., Philips, A., Antel, J., Healy, L., Moore, C.S., Bendotti, C., Aronica, E., Radulescu, C.I., Barnes, S.J., Hampton, D.W., van der Valk, P., Jacobson, S., Matthews, P.M., Amor, S., Owen, D.R., 2022. Translocator protein is a marker of activated microglia in rodent models but not human neurodegenerative diseases. bioRxiv, 2022.2005.2011.491453.

O’Connor, J.C., Lawson, M.A., Andre, C., Moreau, M., Lestage, J., Castanon, N., Kelley, K.W., Dantzer, R., 2009. Lipopolysaccharide-induced depressive-like behavior is mediated by indoleamine 2,3-dioxygenase activation in mice. Mol Psychiatry 14, 511–522.

O’Donnell, L.J., Pasternak, O., 2015. Does diffusion MRI tell us anything about the white matter? An overview of methods and pitfalls. Schizophr Res 161, 133–141.

Pape, K., Tamouza, R., Leboyer, M., Zipp, F., 2019. Immunoneuropsychiatry – novel perspectives on brain disorders. Nat Rev Neurol 15, 317–328.

Papp, E.A., Leergaard, T.B., Calabrese, E., Johnson, G.A., Bjaalie, J.G., 2014. Waxholm Space atlas of the Sprague Dawley rat brain. Neuroimage 97, 374–386.

Pasternak, O., Sochen, N., Gur, Y., Intrator, N., Assaf, Y., 2009. Free water elimination and mapping from diffusion MRI. Magn Reson Med 62, 717–730.

Pasternak, O., Westin, C.F., Bouix, S., Seidman, L.J., Goldstein, J.M., Woo, T.U., Petryshen, T.L., Mesholam-Gately, R.I., McCarley, R.W., Kikinis, R., Shenton, M.E., Kubicki, M., 2012. Excessive extracellular volume reveals a neurodegenerative pattern in schizophrenia onset. J Neurosci 32, 17365–17372.

Polsek, D., Cash, D., Veronese, M., Ilic, K., Wood, T.C., Milosevic, M., Kalanj-Bognar, S., Morrell, M.J., Williams, S.C.R., Gajovic, S., Leschziner, G.D., Mitrecic, D., Rosenzweig, I., 2020. The innate immune toll-like-receptor-2 modulates the depressogenic and anorexiolytic neuroinflammatory response in obstructive sleep apnoea. Sci Rep 10, 11475.

Radhakrishnan, H., Shabestari, S.K., Blurton-Jones, M., Obenaus, A., Stark, C.E.L., 2022. Using Advanced Diffusion-Weighted Imaging to Predict Cell Counts in Gray Matter: Potential and Pitfalls. Front Neurosci 16, 881713.

Schilling, K.G. Grussu, F., Ianus, A., Hansen, B., Aggarwal, M., Michielse, S., Nasrallah, F., Syeda, W., Wang, N., Veraart, J., Roebroeck, A., Bagdaserian, A.F., Eichner, C., Sepehrband, F., Zimmermann, J., Jeurissen, B., Frydman, L., van de Looij, Y., Hike, D., Dunn, J. F., Miller, K. L., Landman, B. A., Shemesh, N., Anderson, A., McKinnon, E., Farquharson, S., Dell’ Acqua, F., Pierpaoli, C., Drobnjak, I., Leemans, A., Harkins, K. D., Descoteaux, M., Xu, D., Santin, M. D., Grant, S., Obenaus, A., Kim, G. S., Wu, D., Le Bihan, D., Blackband, S. J., Ciobanu, L., Fieremans, E., Bai, R., Leergaard, T., Zhang, J., Dyrby, T. B., Johnson, G. A., Cohen-Adad, J., Budde, M. D., Jelescu, I. O., 2022. Recommendations and guidelines from the ISMRM Diffusion Study Group for preclinical diffusion MRI: Part 2 – Ex vivo imaging. p. arXiv:2209.13371.

Sehy, J.V., Ackerman, J.J., Neil, J.J., 2002. Evidence that both fast and slow water ADC components arise from intracellular space. Magn Reson Med 48, 765–770.

Smith, S.M., Jenkinson, M., Woolrich, M.W., Beckmann, C.F., Behrens, T.E., Johansen-Berg, H., Bannister, P.R., De Luca, M., Drobnjak, I., Flitney, D.E., Niazy, R.K., Saunders, J., Vickers, J., Zhang, Y., De Stefano, N., Brady, J.M., Matthews, P.M., 2004. Advances in functional and structural MR image analysis and implementation as FSL. Neuroimage 23 Suppl 1, S208–219.

Stock, A.D., Gelb, S., Pasternak, O., Ben-Zvi, A., Putterman, C., 2017. The blood brain barrier and neuropsychiatric lupus: new perspectives in light of advances in understanding the neuroimmune interface. Autoimmun Rev 16, 612–619.

Stolp, H.B., Ball, G., So, P.W., Tournier, J.D., Jones, M., Thornton, C., Edwards, A.D., 2018. Voxel-wise comparisons of cellular microstructure and diffusion-MRI in mouse hippocampus using 3D Bridging of Optically-clear histology with Neuroimaging Data (3D-BOND). Sci Rep 8, 4011.

Tan, Y.L., Yuan, Y., Tian, L., 2020. Microglial regional heterogeneity and its role in the brain. Mol Psychiatry 25, 351–367.

Tschanz, S.A., Burri, P.H., Weibel, E.R., 2011. A simple tool for stereological assessment of digital images: the STEPanizer. J Microsc 243, 47–59.

Turkheimer, F., Veronese, M., Mondelli, V., Cash, D., Pariante, C., 2022. Sickness Behaviour and Depression: An Updated Model of Peripheral-Central Immunity Interactions. Preprints 2022030062.

Turkheimer, F.E., Althubaity, N., Schubert, J., Nettis, M.A., Cousins, O., Dima, D., Mondelli, V., Bullmore, E.T., Pariante, C., Veronese, M., 2021. Increased serum peripheral C-reactive protein is associated with reduced brain barriers permeability of TSPO radioligands in healthy volunteers and depressed patients: implications for inflammation and depression. Brain Behav Immun 91, 487–497.

Valdes-Hernandez, P.A., Sumiyoshi, A., Nonaka, H., Haga, R., Aubert-Vasquez, E., Ogawa, T., Iturria-Medina, Y., Riera, J.J., Kawashima, R., 2011. An in vivo MRI Template Set for Morphometry, Tissue Segmentation, and fMRI Localization in Rats. Front Neuroinform 5, 26.

Van Camp, N., Lavisse, S., Roost, P., Gubinelli, F., Hillmer, A., Boutin, H., 2021. TSPO imaging in animal models of brain diseases. Eur J Nucl Med Mol Imaging 49, 77–109.

Vicente-Rodriguez, M., Singh, N., Turkheimer, F., Peris-Yague, A., Randall, K., Veronese, M., Simmons, C., Karim Haji-Dheere, A., Bordoloi, J., Sander, K., Awais, R.O., Arstad, E., Nima, C., Cash, D., Parker, C.A., 2021. Resolving the cellular specificity of TSPO imaging in a rat model of peripherally-induced neuroinflammation. Brain Behav Immun 96, 154–167.

West, M.J., Slomianka, L., Gundersen, H.J., 1991. Unbiased stereological estimation of the total number of neurons in thesubdivisions of the rat hippocampus using the optical fractionator. Anat Rec 231, 482–497.

Winkler, A.M., Ridgway, G.R., Webster, M.A., Smith, S.M., Nichols, T.E., 2014. Permutation inference for the general linear model. Neuroimage 92, 381–397.

Yi, S.Y., Barnett, B.R., Torres-Velazquez, M., Zhang, Y., Hurley, S.A., Rowley, P.A., Hernando, D., Yu, J.J., 2019. Detecting Microglial Density With Quantitative Multi-Compartment Diffusion MRI. Front Neurosci 13, 81.

Zhang, H., Schneider, T., Wheeler-Kingshott, C.A., Alexander, D.C., 2012. NODDI: practical in vivo neurite orientation dispersion and density imaging of the human brain. Neuroimage 61, 1000–1016.

Zhang, L., Hu, K., Shao, T., Hou, L., Zhang, S., Ye, W., Josephson, L., Meyer, J.H., Zhang, M.R., Vasdev, N., Wang, J., Xu, H., Wang, L., Liang, S.H., 2021. Recent developments on PET radiotracers for TSPO and their applications in neuroimaging. Acta Pharm Sin B 11, 373–393.

Zhao, J., Bi, W., Xiao, S., Lan, X., Cheng, X., Zhang, J., Lu, D., Wei, W., Wang, Y., Li, H., Fu, Y., Zhu, L., 2019. Neuroinflammation induced by lipopolysaccharide causes cognitive impairment in mice. Sci Rep 9, 5790.

