## Appendix - NIMA Members for "Mapping acute neuroinflammation *in vivo* with diffusion-MRI in rats given a systemic lipopolysaccharide challenge"

**NIMA members and affiliations during Part 1 workpackages (2014-17)**

Brighton & Sussex University Hospitals NHS Trust

Dominika Wlazly

Cambridgeshire & Peterborough NHS Foundation Trust

Amber Dickinson, Andy Foster, Clare Knight

Cardiff University

Claire Leckey, Paul Morgan, Angharad Morgan, Caroline O'Hagan, Samuel Touchard

GSK

Shahid Khan, Phil Murphy, Christine Parker, Jai Patel, Jill Richardson

Janssen

Paul Acton, Nigel Austin, Anindya Bhattacharya, Nick Carruthers, Peter de Boer, Wayne Drevets, John Isaac, Declan Jones, John Kemp, Hartmuth Kolb, Jeff Nye, Gayle Wittenberg

Kings College London

Gareth Barker, Anna Bogdanova, Heidi Byrom, Diana Cash, Annamaria Cattaneo, Daniela Enache, Tony Gee, Caitlin Hastings, Melisa Kose, Giulia Lombardo, Nicole Mariani, Anna McLaughlin, Valeria Mondelli, Maria Nettis, Naghmeh Nikkheslat, Carmine Pariante, Karen Randall, Julia Schubert, Luca Sforzini, Hannah Sheridan, Camilla Simmons, Nisha Singh, Federico Turkheimer, Vicky Van Loo, Mattia Veronese, Marta Vicente Rodriguez, Toby Wood, Courtney Worrell, Zuzanna Zajkowska

Lundbeck

Brian Campbell, Jan Egebjerg, Hans Eriksson, Francois Gastambide, Karen Husted Adams, Ross Jeggo, Thomas Moeller, Bob Nelson, Niels Plath, Christian Thomsen, Jan Torleif Pederson, Stevin Zorn

NHS Greater Glasgow and Clyde

Catherine Deith, Scott Farmer, John McClean, Andrew McPherson, Nagore Penandes, Paul Scouller, Murray Sutherland

Oxford Health NHS Foundation Trust

Mary Jane Attenburrow, Jithen Benjamin, Helen Jones, Fran Mada, Akintayo Oladejo, Katy Smith

Pfizer

Rita Balice-Gordon, Brendon Binneman, James Duerr, Terence Fullerton, Veeru Goli, Zoe Hughes, Justin Piro, Tarek Samad, Jonathan Sporn

Sussex Partnership NHS Foundation Trust

Liz Hoskins, Charmaine Kohn, Lauren Wilcock

University of Cambridge

Franklin Aigbirhio, Junaid Bhatti, Ed Bullmore, Sam Chamberlain, Marta Correia, Anna Crofts, Tim Fryer, Martin Graves, Alex Hatton, Manfred Kitzbichler, Mary-Ellen Lynall, Christina Maurice, Ciara O'Donnell, Linda Pointon, Peter St George Hyslop, Lorinda Turner, Petra Vertes, Barry Widmer, Guy Williams

University of Glasgow

Jonathan Cavanagh, Alison McColl, Robin Shaw

University of Groningen

Erik Boddeke

University of Oxford

Alison Baird, Stuart Clare, Phil Cowen, I-Shu (Dante) Huang, Sam Hurley, Simon Lovestone, Alejo Nevado-Holgado, Elena Ribe, Anviti Vyas, Laura Winchester

University of Southampton

Madeleine Cleal, Diego Gomez-Nicola, Renzo Mancuso, Hugh Perry

University of Sussex

Mara Cercignani, Charlotte Clarke, Alessandro Colasanti, Neil Harrison, Rosemary Murray

University of Texas

Jason O'Connor

University of Toronto

Howard Mount
