## Supplemental Figures for "Mapping acute neuroinflammation *in vivo* with diffusion-MRI in rats given a systemic lipopolysaccharide challenge"

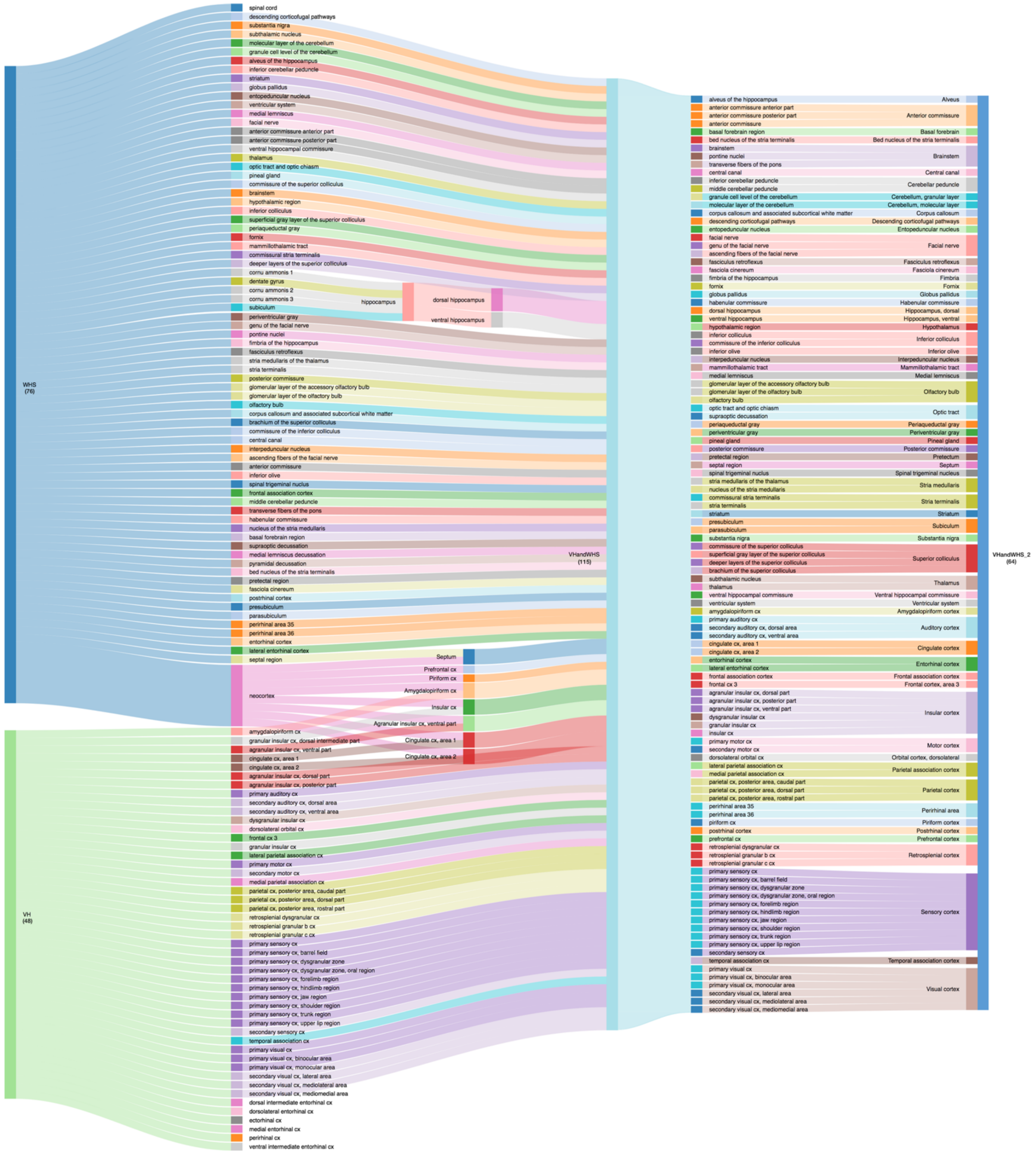


Supplemental Figure 1 A Sankey diagram showing how regions of interest (ROI) from the Waxholm Space (WHS) atlas (76 ROIs) and the Valdes-Hernandez (VH) atlas (48 ROIs) were combined, split, and merged to create the VHandWHS atlas (115 ROIs) and the VHandWHS_2 atlas (64 ROIs). These 64 ROIs were used for ROI-based analyses in this study. Most of the neocortex ROI from the WHS atlas overlapped with and was replaced by ROIs from the VH atlas. The remainder of the neocortex ROI was split as shown in the diagram. The ectorhinal and entorhinal cortex ROIs from the VH atlas overlapped with and were replaced by the perirhinal and entorhinal cortex ROIs from the WHS atlas. The spinal cord ROI from the WHS atlas, considered a region of no interest, was removed.

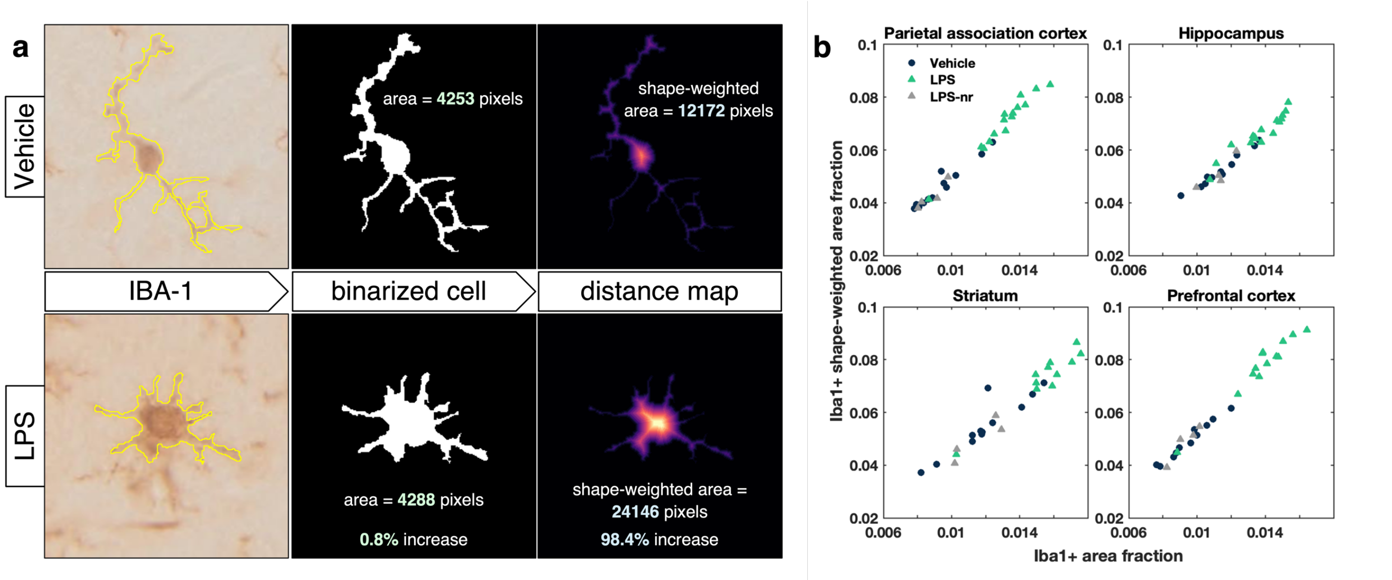


Supplemental Figure 2 Shape-weighted area fraction (SWAF) vs. simple area fraction for assessing microglial morphology. a) An illustration of the calculation of SWAF for microglia from a vehicle-treated brain (top) and an LPS-treated brain (bottom). The microglia have nearly identical areas (number of pixels), but the shape-weighted area (sum of values in the intracellular distance map) of the reactive microglia is nearly twice that of the non-reactive microglia. b) Scatter plots showing strong correlations between the SWAF and simple area fraction. Each symbol represents an individual rat. Blue circles represent control rats, green triangles LPS rats, and gray triangles LPS non-responders.

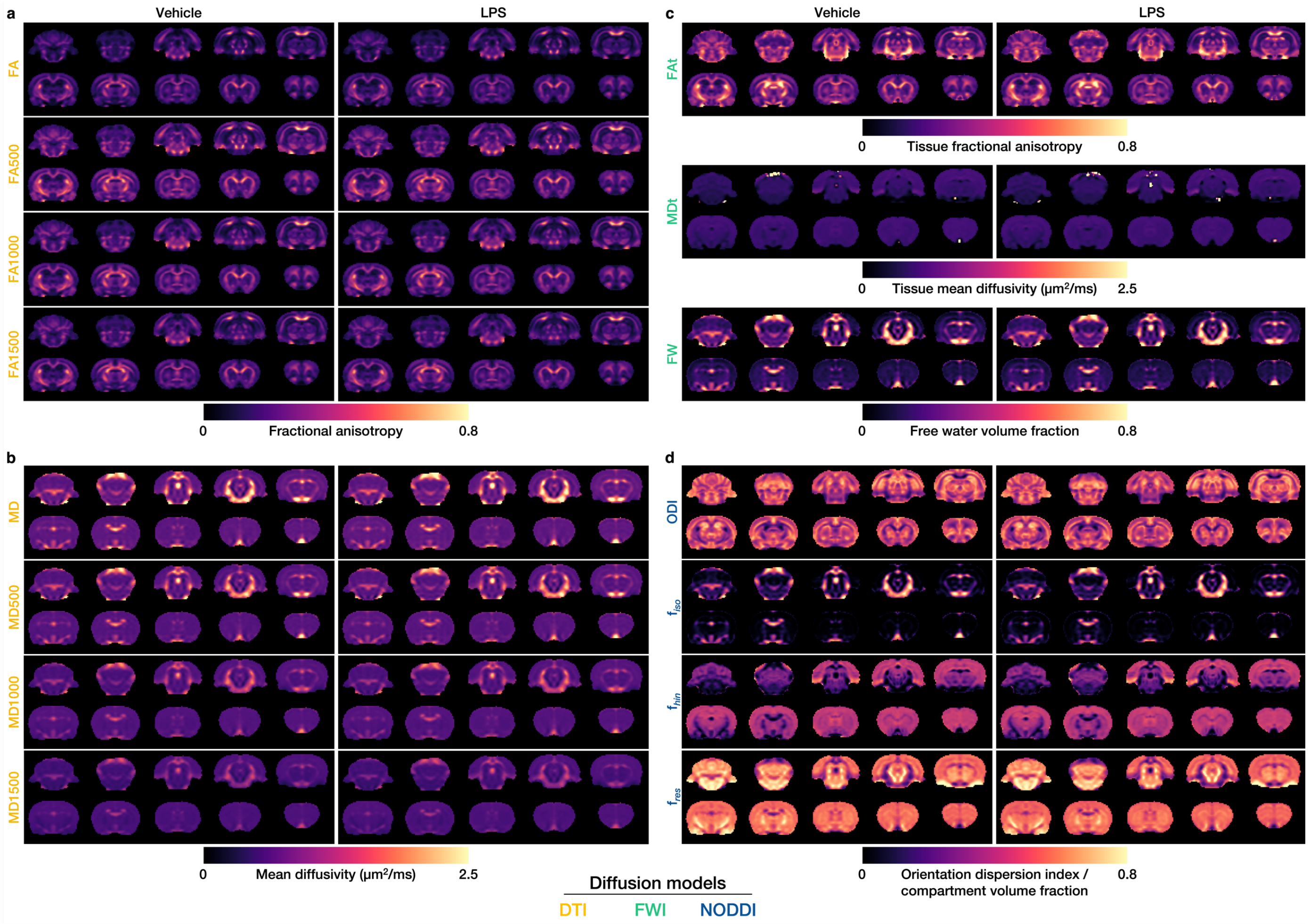


Supplemental Figure 3 Group mean maps of parameters from three different diffusion models. a) Fractional anisotropy from the simple diffusion tensor imaging (DTI) model using all three shells (FA), just the b500 shell (FA500), just the b1000 shell (FA1000), and just the b1500 shell (FA1500). b) Mean diffusivity from the simple DTI model using all three shells (MD), just the b500 shell (MD500), just the b1000 shell (MD1000), and just the b1500 shell (MD1500). c) Tissue fractional anisotropy (FAt), tissue mean diffusivity (MDt), and free water volume fraction (FW) from the free water imaging (FWI) model. d) Orientation dispersion index (ODI), isotropic diffusion volume fraction ($f_{iso}$), hindered diffusion volume fraction ($f_{hin}$), and restricted diffusion volume fraction ($f_{res}$) from the neurite orientation dispersion and density imaging (NODDI) model.

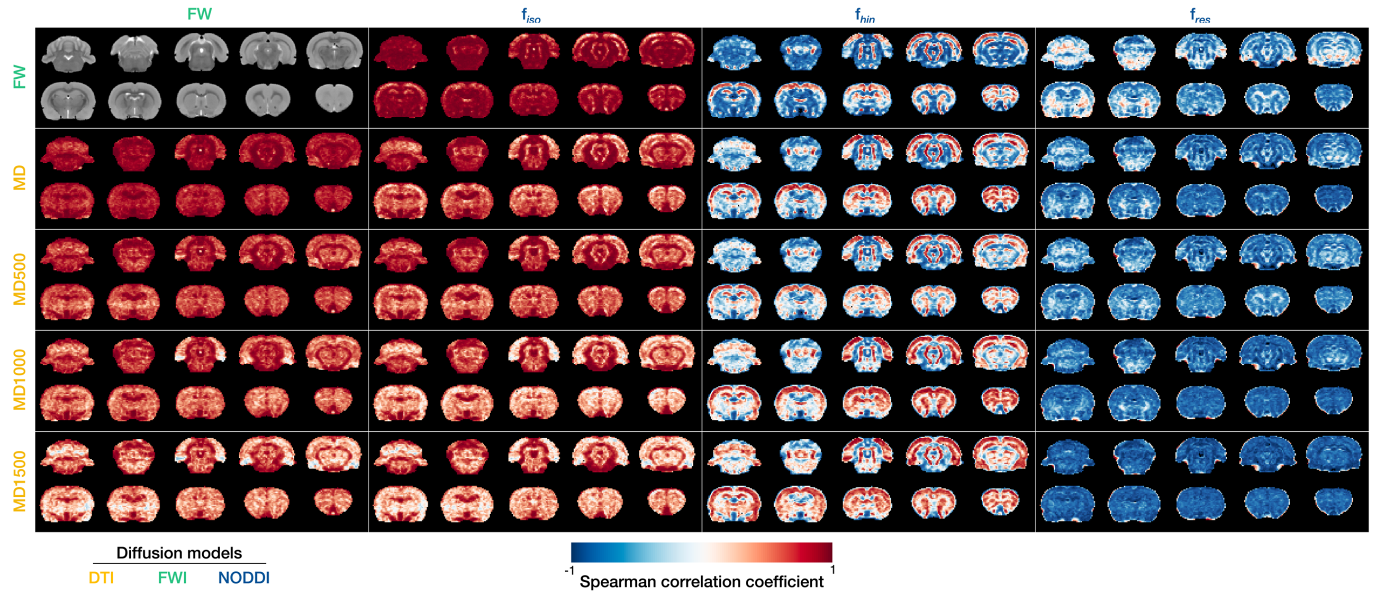


Supplemental Figure 4 Voxel-wise Spearman correlation coefficients across rats between parameters from three different diffusion models. Rows, top to bottom: free water volume fraction (FW) from the free water imaging (FWI) model; mean diffusivity from the simple diffusion tensor imaging (DTI) model using all three shells (MD), just the b500 shell (MD500), just the b1000 shell (MD1000), and just the b1500 shell (MD1500). Columns, left to right: FW from the FWI model; and isotropic diffusion volume fraction ($f_{iso}$), hindered diffusion volume fraction ($f_{hin}$), and restricted diffusion volume fraction ($f_{res}$) from the neurite orientation dispersion and density imaging (NODDI) model. Coronal slices of a T2-weighted rat brain template are shown in the top left corner for reference.

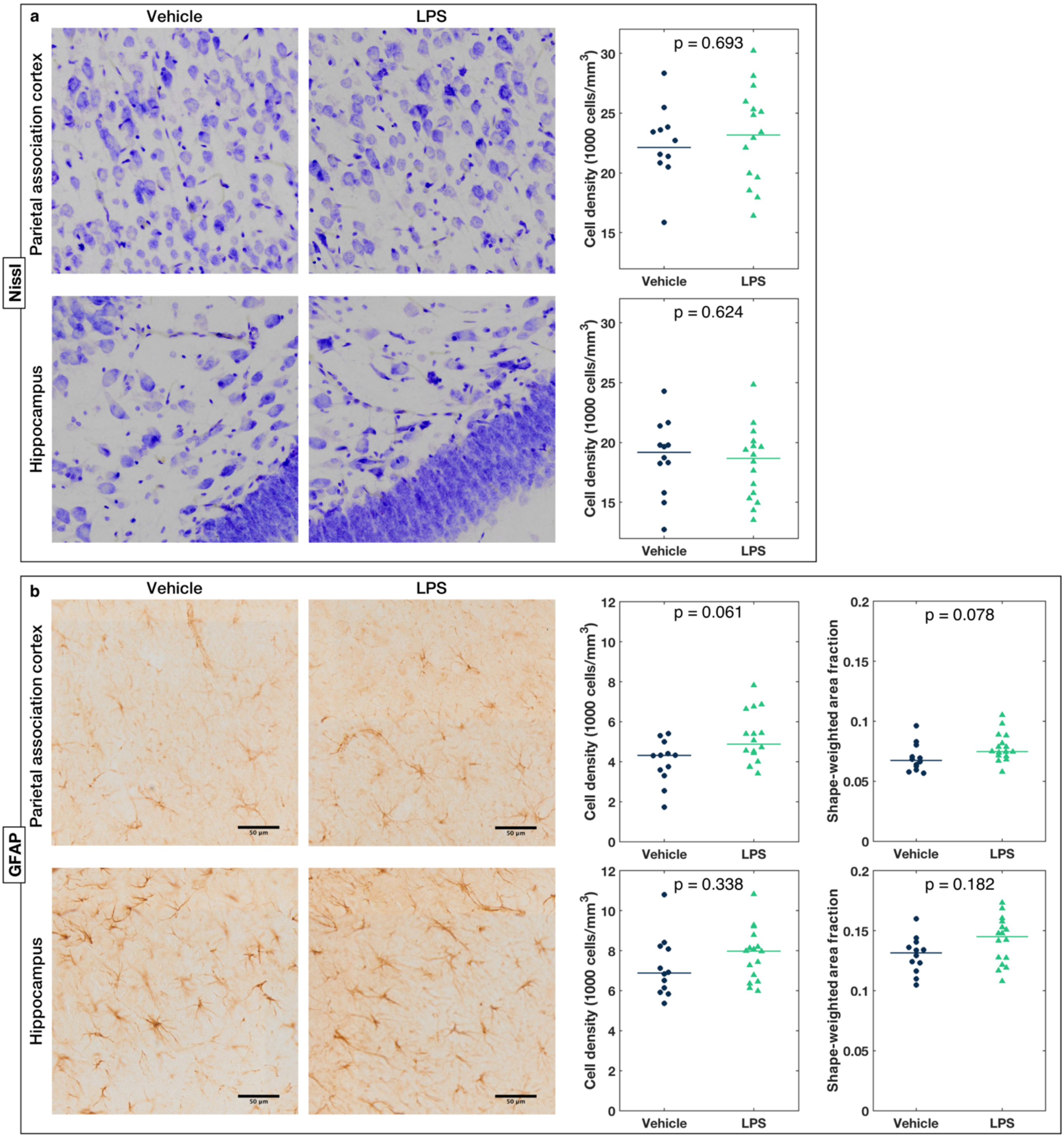


Supplemental Figure 5 Representative 40× images of a) Nissl- and b) GFAP-stained brain sections from the parietal association cortex and hippocampus of a vehicle-treated control (first column) and LPS-treated rat (second column). Third column: Dot plots of stereologically estimated a) Nissl+ and b) GFAP+ cell density in the parietal association cortex and hippocampus of individual rats. Fourth column: b) Dot plots of shape-weighted area fraction of GFAP+ cells in the parietal association cortex and hippocampus of individual rats. Horizontal lines indicate group medians. P-values are from two-tailed Mann-Whitney U tests and FDR-corrected.

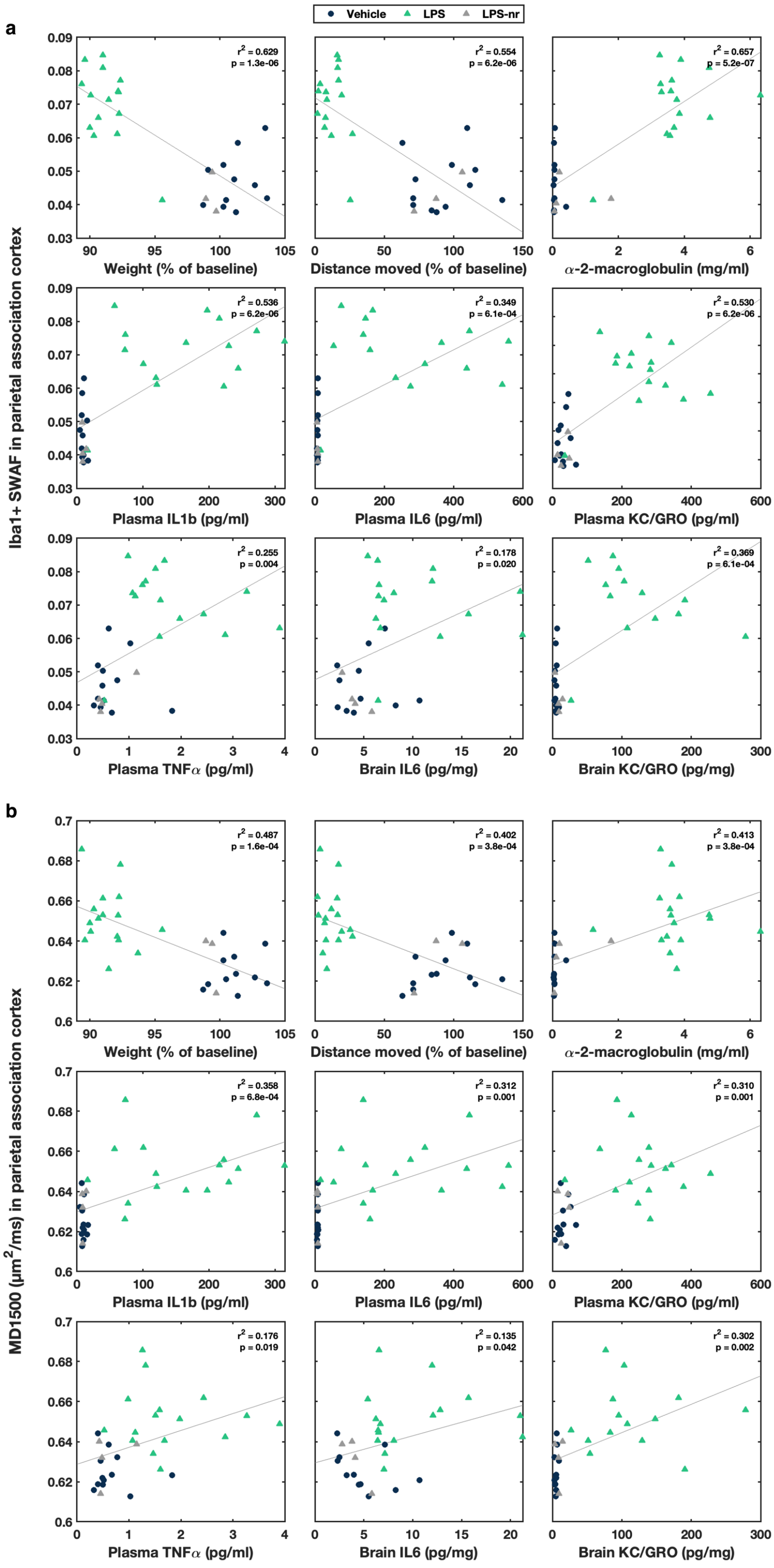


Supplemental Figure 6 a) Histologically measured Iba1+ shape-weighted area fraction (SWAF) and b) median values of b1500 mean diffusivity (MD1500) in the parietal association cortex vs. non-imaging measures. Each symbol represents an individual rat. Blue circles represent control rats, green triangles LPS rats, and gray triangles LPS non-responders. Linear regression lines (all rats pooled) are plotted in gray, and the R-squared statistic and uncorrected p-value of each regression model displayed in each plot.
